# Adaptive Mating Among Natural Strains of *S. cerevisiae*

**DOI:** 10.1101/2024.02.18.580925

**Authors:** Sivan Kaminski Strauss, Gianni Liti, Orna Dahan, Yitzhak Pilpel

## Abstract

In evolutionary biology, sexual mating plays a pivotal role in facilitating the combination of beneficial alleles among individuals. Cross-species data suggest that organisms selectively mate with partners based on factors such as genetic distance and partner fitness. Understanding the determinants of pair-specific mating affinity is crucial for unraveling the impact of sex on evolution. However, despite the significance of this phenomenon, the availability of large and consistent datasets is limited, leading to inconsistent conclusions.

To address this gap, we present a comprehensive mating assay enabling the simultaneous quantification of mating affinity among approximately 100 natural *Saccharomyces cerevisiae* strains. Our study demonstrates that mating in an *en masse* manner, allowing mating based on affinity, enhances the overall fitness of the hybrids offspring population compared to mating that enforces one specific partner at a time. By employing a DNA barcode recombination system integrated into natural isolates’ genomes, we sequence recombined barcode pairs and revealed mating frequencies and affinities among all strains in different environments.

Our findings unveil strain-specific mating affinity among natural yeast strains, with certain parental pairs exhibiting a heightened affinity for each other over other strains, whereas certain strains combinations are avoided. Notably, among the pairs with the highest affinity, there is a preference for lower genetic distances. Intriguingly, multiple strains show a propensity for mating with partners that yield higher-fitness hybrids on average. Collectively, our results provide compelling evidence that yeast actively engages in adaptive mate affinity.

## Introduction

Mate selection is a widespread phenomenon in nature, observed across various species, including fungi, yeast, plants, insects, fish, lizards, amphibians, birds, and mammals [1]–[18]. The affinity between mating partners, a combination of selection and sexual compatibility, can be either pair-specific or general, where certain individuals may exhibit a higher attractiveness level than others. Assortative mating, a pivotal mechanism in achieving pair-specific affinity, involves individuals with similar phenotypes or genotypes exhibiting a higher mating frequency. Extensive study of this phenomenon and its converse, i.e. disassortative mating, has been conducted across diverse organisms [19]–[21].

In certain species, mate selection has been shown to confer advantages, yielding highly fit offspring [22]– [24]. Notably, the avoidance of partners with high genetic similarity, disassortative mating, has been observed in various organisms and is also correlated with higher-fitness offspring [25]–[29]. In contrast, mating with partners with low genetic similarity has been linked to the production of sterile or low-fit offspring [29]–[32]. Consequently, maximizing offspring fitness involves selecting mates within an intermediate range of genetic similarity between parents [8]. Recent studies, including those on yeast, plants, and mice, indicate an optimal genetic distance that maximizes offspring fitness [33].

The budding yeast, *Saccharomyces cerevisiae*, grows either vegetatively as haploid or as diploid and it can also undergo sexual mating. Pheromone sensing initiates the mating process, resulting in haploid cells (Mat**a** and Matα) forming diploid cells that continue growing vegetatively, referred to here as “hybrids” or “hybrids offspring”. Diploid cells undergo sporulation and meiosis upon nitrogen- and carbon-starvation, producing haploid spores stored within the mother cell’s intact cytoplasm, known as the ascus [34]. While yeast predominantly mate with sibling haploid cells near spore germination [35], [36], previous studies indicate occurrences of outcrossing and haplo-selfing, albeit relatively infrequently [37], influencing genome mixing through recombination [38]–[45].

As outcrossing plays a vital role in shaping yeast population structure, an unanswered question pertains to the extent of pair-specific mating affinity exerted among yeast strains. Historically, it has been established that a crucial factor influencing mating affinity in *S. cerevisiae* is the level of pheromone secretion, leading to a preference for mating with the partners exhibiting the highest pheromone-secreting capability [46]–[49]. These key studies were done mainly by genetically manipulating pheromone levels and other mating related genes in a handful of strains and examining the effect on mate affinity. More recently, another criterion has been demonstrated to impact affinity in yeast mating, the cell size. In conditions where cell size is crucial for cell fitness, haploid cells tend to choose partners whose cell size aligns more closely with the optimal size for fitness in that particular condition [24]. A recent study [50] showed that a single point mutation that changes a tryptophan amino acid to arginine in the GPA1 gene reduces fitness of the strains, but increases their mating efficiency. Moreover, this mutation was also associated with increased outbreeding tendency of natural strains. However, the influence of additional factors such as overall fitness of either the mating partners (or perhaps even the resulting diploids) and genetic similarity between Mat**a** and Matα haploids on mating affinity between yeast pairs remains to be fully elucidated. Furthermore, the mating patterns in natural isolates was never explored at a high-throughput manner to our knowledge.

To explore the extent mating affinity in yeast and the factors that govern it, we employed a high-throughput approach, utilizing a collection of unprecedentedly large natural *S. cerevisiae* strains. We employed a novel mating-recording barcode strategy in a pooled mating assay with ∼100 natural strains that enabled the tracking of thousands of hybrids offspring. In a recent paper [51], we used this system to explore the modes of fitness inheritance in yeast. We found that fitness inheritance mode vary between the two main lifestyles of yeast, fermentation and respiration; hybrids fitness positively correlated with the fitness of its parents in fermentation and with the genetic distance between its parents in respiration [51]. Here, we turn to track, using this barcode recombination system the rate of formation of thousands of hybrid offspring, offering a unique opportunity to explore mating dynamics in an organism in unprecedented extent and natural diversity. We observe that selective mate affinity among strains significantly enhances the fitness of the resultant hybrid offspring population with a higher affinity towards partners yielding higher average fitness. Additionally, we observe dynamic shifts in mating affinity towards different strain patterns under varying environmental conditions. Finally, our study reveals higher affinity for strains with short to intermediate genetic distances.

## Results

### Partner exploration during mating enhances offspring fitness

To study mating affinity in yeast, and to examine its effects on hybrids’ fitness, we developed a genetic system as described in Strauss *et al* [51]. In short, we used a pool of ∼100 *S. cerevisiae* natural isolates selected from a world-wide collection that represents the species diversity (taken from [40]). The 100 strains were chosen to vary in their fitness on fermentative and respiratory media types and to span a broad range of pairwise genetic distances (GD) (**Figure 1A**).

**Figure 1.**
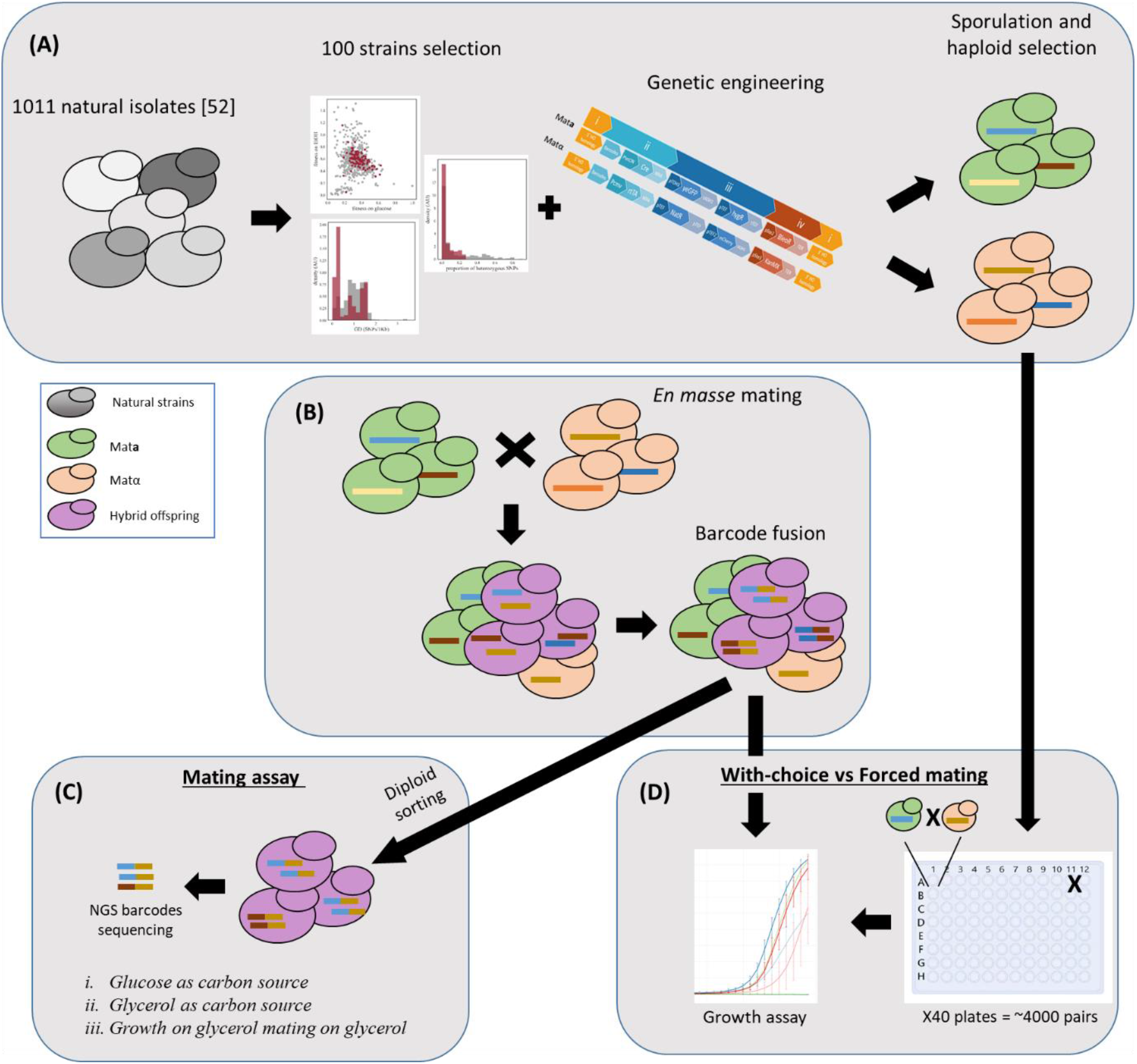
The mating assay design. (A) 100 strains from Peter *et al.* [40] 1011 strains were chosen according to three parameters; their fitness under respiration and fermentation conditions, the genetic distance (GD) of strain pairs in the collection, and heterozygosity levels. The 100 strains were genetically manipulated to contain fluorescence and antibiotic markers and a barcode fusion genetic system. Mat**a** and Matα final strains were achieved by sporulation and haploid selection (for more information regarding strain choice and genomic engineering, see Strauss *et al*. [51]) (B) All strains from both mating types (Mat**a** and Matα) were mixed together and allowed to mate with one another, *en masse*. Barcode recombination was enabled during mating. After mating, culture was sorted for hybrid offspring cells (based on expressing the two florescent markers) to measure pairing (C) or used as a pool for growth measurements (D) (C) Mating assay: after sorting, to recover hybrid offspring only, fused barcodes were sequenced to reveal parents’ identity of each strain. Mating assays were done on three conditions; (i) Glucose or (ii) Glycerol as carbon sources and (iii) strains were grown on Glucose and switched to Glycerol for mating (D) All individual parents were used for both *en masse* mating (“With choice”) and for pairwise mating (“Forced”). Following mating both populations were grown and OD was measured

To assess the direct impact of mating affinity on hybrids’ fitness, we conducted a comparative analysis of the growth rates between two distinct offspring populations: one with the ability to explore mating partners (’With-choice’) and another without this ability (’Forced’). The ’With-choice’ hybrids pool was generated by combining all haploid strains and allowing them to mate *en masse* based on their mutual ’affinities’. In contrast, the ’Forced’ hybrids population resulted from individually conducting all possible pairwise mating, employing 96-well plates, and subsequently pooling the offspring (**Figure 1B and 1D**). Mating were conducted using either glucose or glycerol as a carbon source that represent the two major energy metabolism in yeast fermentation and aerobic respiration respectively. We then measured the growth rates of the two population pools under the two respective conditions, glucose or glycerol. To ensure accurate growth measurements of the hybrids offspring, we inhibit haploid growth by introducing two antibiotics that inhibits the growth of either Mat**a** or Matα and included vigorous shaking during the incubation periods to prevent further mating.

**Figure 2A** shows that, under both fermentative and respiratory conditions, the ’With-choice’ population, on each of three biological repeats, exhibited superior growth dynamics compared to the ’Forced’ pool. Notably, in the respiratory condition, a very pronounced difference between the two pools was observed. The hybrids pool resulting from the ’With-choice’ mating displayed much accelerated growth (i.e., faster growth rate) and a higher yield (i.e final cell density at the stationary phase) compared to the ’Forced’ populations. In the glucose condition, the advantage of the ’With-choice’ pool was more subtle, manifested through a higher growth rate during the logarithmic phase and enhanced performance after the diauxic shift regime in which yeast cells switch from fermentation to aerobic respiration (approximately observed from 15 hours onward). This experiment provides compelling evidence that affinities towards specific mating partners in yeast significantly enhances the fitness of hybrid offspring, with a particularly pronounced effect in respiratory medium conditions.

**Figure 2.**
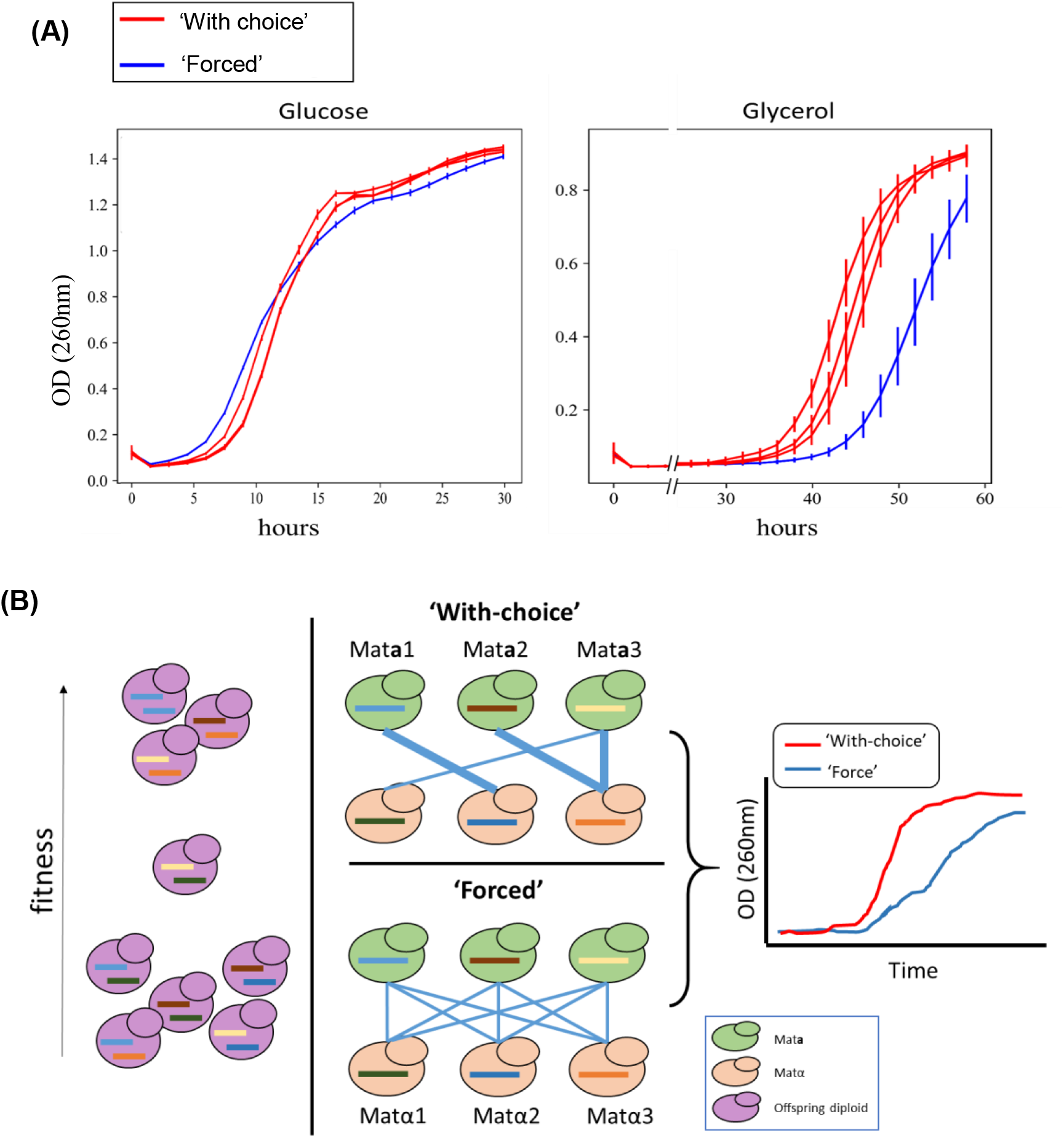
Pool of hybrids produced with “choice” have higher fitness than offspring produced with no such choice, especially in the respiratory conditions. (A) Left: glucose as carbon source, right: glycerol as carbon source. OD600 was measured every two hours for 35 hours (left) or 60 hours (right). Each curve is the mean of 14 technical repeats and error bars represents the standard deviation of the 14 repeats Red lines are “With choice” mating (3 biological repeats), blue line is the “Forced” mating. (B) Hypothetical explanation to the observation in (A). Assuming hybrids’ fitness ranks are as in the left panel (i.e. the 3 hybrid types on top having the highest fitness, and the 5 at the bottom of the figure having the lowest). In the middle, bottom panel: all offspring combinations are forced to be made in similar amounts. Middle, upper panel: In the ‘With-choice’ mating, parents can explore partners and mate based on mating affinity potentially producing more high-fit hybrids than in the ‘Forced’ option where each parent mixed with each partners separately. As such, the population made from the ‘With-choice’ mating is biased toward high fitness hybrids, as can be detected by measuring the growth curves of the population (right panel)

A plausible explanation is that while in the ’Forced’ population all combinations of offspring were made (based on mating efficiency between parental strains only), in the ’With-choice’ population, parents could express affinities to specific partners with which hybrids’ fitness is high and thus biased the hybrid population toward a high fitness population (**Figure 2B**)

### Mating assay reveals pair-specific avoidance

To precisely track the mating pairing of each strain during mating in the ’With-choice’ mode, we employed a barcod recombination system enabling the quantification of mating interactions between all pairs of strains (**Figure 1A** and [51]). In brief, we disrupted the HO locus [52], [53] using a construct containing an antibiotic resistance cassette, a fluorescence marker, and a barcode recombination element based on the Cre-lox recombinase system (barcode fusion genetics, BFG) [54] (and similarly to [55]) (see **Figure S1**). This cassette facilitates the recombination of unique haploid parental barcodes from the two mating partners, Mat**a** and Matα into a fused barcode within the same DNA segment, allowing subsequent parental identification. A comprehensive description of this system can be found in [51]. As described above, mating experiments were conducted under two distinct nutritional conditions: a fermentable carbon source, glucose, and a non-fermentable carbon source, glycerol. An additional experimental condition involved growing haploid parental strains on glucose and switching to glycerol immediately before mating was done. In this additional “switching” experiment, we wanted to explore whether strains pairing and affinity is based on the current condition or recent previous conditions that could have influenced their fitness and mutual affinity (**Figure 1B**). Each condition was replicated twice. Our objective was to precisely quantify the frequency at which each potential hybrid is formed. Given that our initial pool comprised 89 parents from the Mat**a** mating type and 46 parents from the Matα mating type, a theoretical maximum of 4,094 potential hybrid combinations could arise. However, our results demonstrated that while 2,681 (approximately 65.3%) hybrid combinations were observed in at least one replicate and under at least one of the conditions investigated, the remaining combinations were never detected. Hybrids observed in different conditions were found to be either exclusive to one condition, present in two, or in all three (**Figure S2A**).

In **Figure 3A**, we depict a matrix with all pairwise combinations, color-coded based on the number of conditions in which they were observed. The color gradient spans from white, indicating no hybrid in any condition, to black, signifying hybrid present in all three tested conditions. The clustering of haploid strains revealed two discernible behaviors among parental pairs with no offspring: one group comprised of parents exhibiting little to no mating activity with any partner across any condition, while the second group consisted of parents actively engaging with many partners but specifically avoiding specific combinations. Additionally, specific strains demonstrated a condition-specific absence of mating activity, where they lacked partners only under a particular condition (**Figure S2B**). Notably, six Matα strains and one Mat**a** strain showed no offspring with any partners across all conditions, including fitness measurement experiments conducted in [51] and their fitness values were incorporated into this study. The hypothetical offspring from these strains (n=574) were excluded from further analysis.

**Figure 3.**
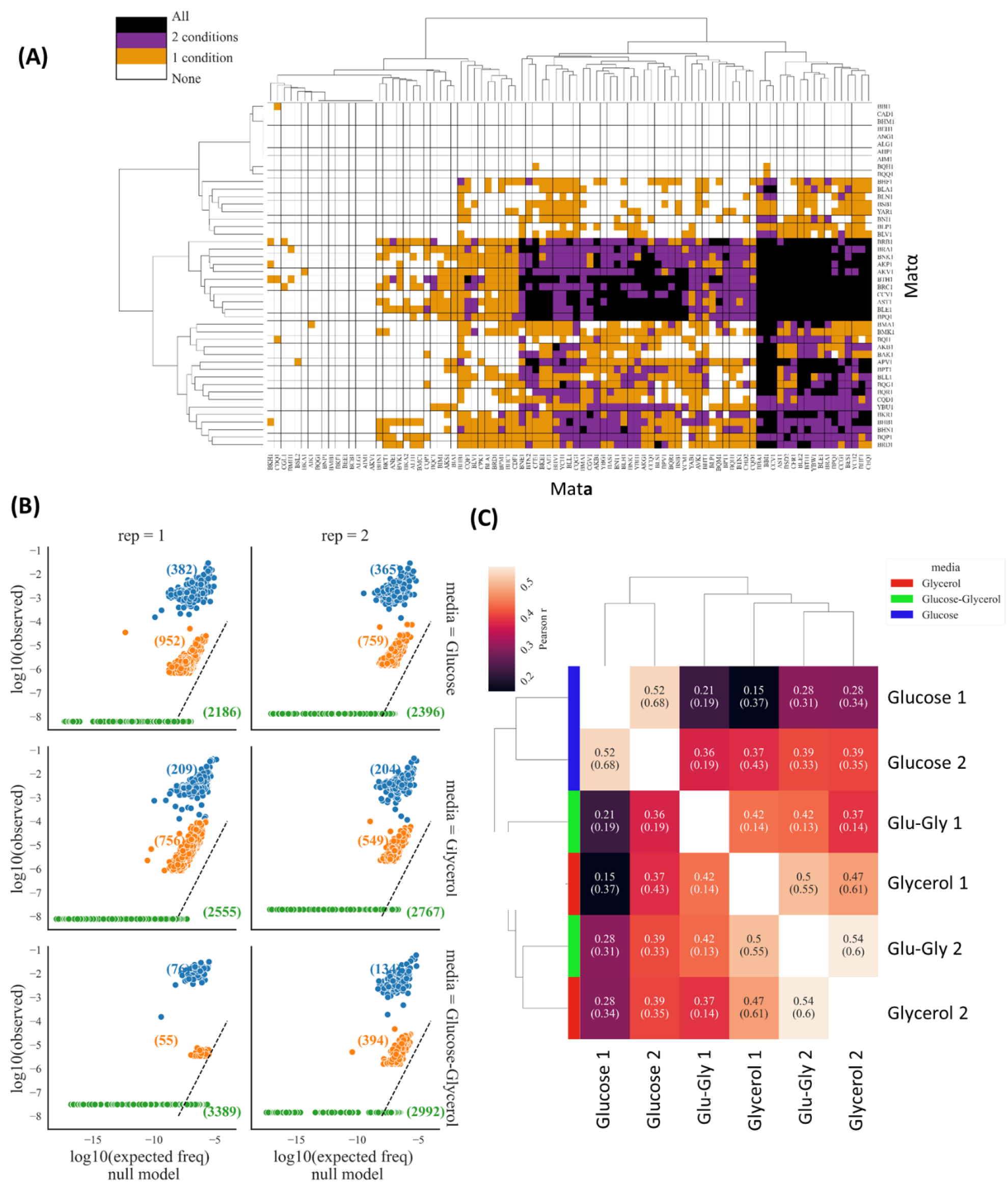
Mating assays reveals enriched and avoided partners. (A) Pair-specific and strain-specific avoidance in *en mass* mating. Each row represents a specific Matα parent, each column represents a specific Mat**a** parents. Each entry in the matrix represents in how many conditions an hybrid of each pair of parents was observed. White represents hybrids that are not detected in any repeat in any condition, black represents hybrids that are being observed in all conditions. Clustering, groups together strains with similar mating patterns, with a dendrogram to the left showing clustering of Mat**α** parents, and a dendrogram above showing the clustering of Mat**a** parents. Pair-specific avoidance is demonstrated as isolated white squares while strain-specific lack of mating activity as white rows or columns. (B) Observed vs expected number of appearances of each hybrid in each condition. Each dot represents a Mat**a**-Matα pair. The y-axis represents the number of observed appearances of the hybrids in the sequencing. The x-axis represents the expected number of appearances based on the product of mating activity of its two parents across all potential partners and of their initial frequencies in the experiment (i.e., null model). Experiments were done in three types of media, and in two repeats from each. Top – glucose, middle – glycerol, bottom – glucose-to-glycerol upon mating; left and right represents the two repetitions. Blue – over-represented hybrids, orange – as expected hybrids, green – absent hybrids. Next to each cluster, in brackets, are the number of hybrid offspring observed. (C) Cluster analysis of mating pairs across conditions. Pearson correlation between the six samples (three conditions and two repeats in each) was calculated based on the observed hybrid frequency normalized to the null model. Hierarchical clustering was done on the matrix of correlation coefficients. The two repeats of glucose (blue in the left column) cluster together, the two repeats of glycerol (green in the left column) cluster with the repeats of the glucose-to-glycerol condition (red in the left column). Pearson r values are denoted in each cell (all p-value are smaller than 10^-20^).

### Selected hybrids in each condition are made either more than, less than, or as expected from their parents’ over-all mating activity

To quantify the amount of hybrids from each pair of parents, we sequenced recombined barcodes from hybrids offspring following their parents *en masse* mating. A notable observation is that the abundance of offspring of various parental pairings spans a broad range, with some being 1,000 more frequently than others, while some pairs not seen at all (**Figure 3B**). We thus hypothesized that this observation represents enhanced or diminished mating affinity. We define mating affinity as the result of a composite, potentially multiplicative effect of efficiency, compatibility, and preference of different strains towards one another. Yet, amounts of hybrids from a pair of parents might also be affect by non-pairwise specific factors. If a Mat**a** parent of a certain strain and a Matα parent from a certain strain exhibit a generally high sexual activity across partners from many strains (e.g. due to high pheromone or receptor levels, or even just due to high amount), they would engage with multiple strains indiscriminately. We would then anticipate, even under a null model assuming no specific affinity between pairs, the production of many hybrids between such generally active strains. Conversely, if two parents are not very active over most partners, we would expect, under a null model of lack of specific affinity, that they might not produce many hybrids between them. We thus constructed a null model that compares and contrasts the observed number of hybrids from each parental pair to the expectation under assumption in which mating features no affinity or repulsion between specific pair of parents (**Figure S3**). This null model was constructed as follows: In each condition, for each pair of parents, we measured the overall mating activity for each of the Mat**a** and Matα parents, as the sum of number of hybrids it had across all potential mating partners in the experiment. Given two parental strains of opposing mating types, the product of their mating activity and their initial frequencies, represents the expected number of hybrids from each pair under the null model (as described in the equation below):

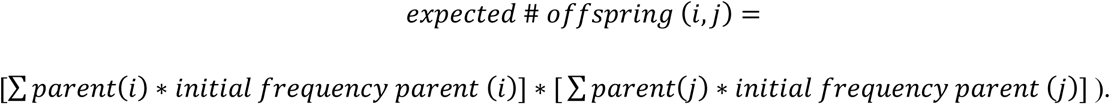

The null model that uses the product of the overall partners of each parent controls for multiple potential confounders. The model considers the background mating activity of each strain, it also allows to eliminate from the calculation other biases that are due to strain-specific mating activity differences, such as pheromone secretion levels, stronger agglutinins, etc.

**Figure 3B** shows the observed number of hybrids of each type vs. the expected number under the null model in each repeat in each of the three conditions. We discern three mating groups defined by observed-to-expected under null model hybrids abundance. The over-represented mating group consists of hybrids that are found to be enriched relative to the null model (**Figure 3B, blue dots**). In the middle, close to the x = y line, is the mating group of hybrids that are observed close to their expected level of abundance, thus reflecting no negative or positive mating selection (**Figure 3B, orange dots**). The lower mating group consists of hybrids that are absent, despite the fact that under the null model many of them should have been observed (**Figure 3B, green dots**). This figure shows that yeast exhibit pair-specific mating affinity; the over represented group consists of partners that mate more with one another, while the absent group reveals pair-specific lack of compatibility or selective avoidance. There are 574 hybrids that were not formed at all in any repeat on any condition. These missing hybrids would have originated from parents that have no hybrids with any other parental strain. These hybrids were removed from this plot.

To further disentangle BFG and mating efficiency from mate affinity, we used pairwise mating and BFG experiments were specific pairs were used to assess BFG and mating efficiencies. We found that BFG efficiency and mating efficiency between individual pairs of strains were not correlated to the mating cluster affiliations (**Figure S4, S6, S5, S7** respectively and **supplementary note**). This result indicates that belonging to the three mate groups does not trivially represent mating or BFG efficiencies between individual strains, but rather points to mate affinity. In addition, we argue that over- or under representation of certain hybrids cannot result, for its most part, from differences in their growth rate post-mating. To limit or even rule out such putative fitness affects, mating was done in conditions that allow for maximum of 4 generations, a very limited time period in which major differences in abundance cannot ensue (see **supplementary note)**

To investigate the potential influence of growth conditions on mating affinity, we clustered the six samples from the two repeats in each of the three conditions. The clustering was based on the correlation between the appearance of hybrids in the six experiments compared to the null model (see **Figure 3C** legend for details). Notably, the two repeats of the glucose condition clustered together; the two repeats of glycerol were co-clustered with the two repeats of the glucose-switched-to-glycerol condition **(Figure 3C**). These results indicate that mating affinity vary in different carbon sources. The fact that the switched condition clustered with the glycerol and not the glucose indicates that mating affinity reflects mainly the prevailing condition in which mating was done rather than the recent growth history of the strains that prevailed immediately until mating.

Recently, it was shown that a single point mutation in the *GPA1* gene (W82R) reduces fitness and induces a higher mating efficiency [50]. We found that 6 Mat**a** strains (YAB1, YCH1, YCH2, AST1, CGL1, CPK1) and 1 Matα (AST) strain from our collection have this mutation. We found moderate, but statistically significant enrichment for those strains’ hybrid offspring to be present in the over-represented mating group in the two glucose repeats (∼1.2-fold increase, p-value<0.01), and in 1 of the glycerol (1.5-fold increase, p-value<0.01) and 1 of the glucose-switched-to-glycerol condition (1.5-fold increase, p-value<0.01). This rather anecdotal observation suggests how a genetic basis might be found for diverse mating behaviors.

### Hybrid fitness and genetic distance modulate mating affinity

We next turned to ask what characterizes hybrids in the over-represented and in the absent groups. As suggested in **Figure 2**, mate choice may favor high-fitness offspring. In order to explore this phenomenon in our system, we partitioned hybrids offspring into four bins based on their relative fitness as measured separately in each growth condition, in Strauss *et al* [51], and examined the fitness distribution in each mating group. We observed a statistically significant enrichment of the highest fitness hybrids in the over-represented mating group across the two repeats of the glucose condition. Conversely, these hybrids were significantly depleted from the under-represented mating group in glucose (**Figure S8A**). In glycerol, the over represented group is enriched with very low fitness hybrids, as indicated by their extinction during hybrids competition experiment (see Strauss *et al*. [51]). This result suggests that in glucose, but not in glycerol, yeast mate more with partners that would on average yield hybrids with high fitness (**Figure S8A**).

We next examined whether the genetic distance between parents differs between the three mating groups. We partitioned parental pairs into four bins of genetic distances between them (as calculated by Peter *et al.* [40] and partitioned in Strauss *et al*. [51]). In the majority of the experiments, we find that parental pairs in the second shortest genetic distance (0.05-0.5 SNPs per 1,000 base pairs) are enriched among the over-represented mating group. This result shows that mating affinity is maximized in this intermediate genetic distance regime (**Figure S8B**). Considering the natural variation in heterozygosity among the natural strains, we hypothesized that either low or high levels of heterozygosity in strains can genetically influence the rate of either inbreeding or outcrossing. We thus hypothesized that such tendencies might manifest in our current mating assay. We indeed find that the enrichment of short-medium genetic distance seen in the over-represented mating group is due to the low heterozygosity strains (<0.08 heterozygous SNPs out of all all SNPs) (**Figure S9**). In contrast, for natural strains with high heterozygosity, no clear and significant trend emerged. These results suggest that strains that did not outcross in their past natural history (low heterozygosity) tend to mate with genetically similar mates, suggesting a genetic determinant for in- or out-crossing.

We next turned to ask if parental strains with high fitness are enriched in the over-represented mating group. We partitioned parents (either Mat**a** or Matα) into four bins based on their relative fitness as measured by competing all Mat**a** and Matα separately on the relevant media (either glucose or glycerol) (Strauss *et al*. [51]). We examined the representation of each of the four fitness bins in each of the three mating groups. We find that in the glucose condition yeast mate more with high fitness partners, while in glycerol they mate more with low fitness partners (**Figure S10**).

To conclude, by harnessing the power of yeast natural isolates collection, we were able to show that yeast mating is influenced by several parameters including the expected hybrids fitness and genetic distance between parents and that this selection depends on the mating conditions and increases the overall fitness of the population.

## Discussion

We have shown, for the first time in a high-throughput experiment, that yeast exhibit pair-specific mating affinity (**Figure 3B**). We further show preferred affinity towards partners with which yeast will tend to have, on average, higher fitness hybrids (**Figure S8A**). In both conditions tested in this paper, yeast mate more with partners that are genetically similar to them, yet not exactly from the same strain (**Figure S8B**), featuring an optimal distance. Additionally, we observed a correlation in which natural strains with low heterozygosity exhibit more frequent mating with partners that are closer genetically to them (**Figure S9**). This finding suggests an inherent tendency of strains to engage in in-crossing or out-crossing.

Along this line, previous research on haplo-selfing in yeast has demonstrated inherent variation among different natural strain isolates regarding their inclination towards engaging in mating after spore germination, a preference which would enhance the probability of inbreeding. This previous paper showed that while some strains have mated in-ascus immediately after spore germinations, others have formed micro-colonies and mated later (i.e., haplo-selfing [43]), supporting the notion that complex mating affinity and behaviors in yeast can be genetically encoded. Apart from genetic distance, in the glucose condition, yeast mate more with high fitness partners, while in glycerol they mate more with low fitness partners (**Figure S10**).

Research has continuously demonstrated the significant role of pheromone secretion levels in the process of mate selection and affinity [46]–[49]. However, the pair-specific mating affinity observed in this study cannot be solely attributed to pheromone levels. Previous findings indicated that cells with higher pheromone secretion are more likely to be selected as mating partners. Yet, this tendency by itself is likely exerted over all potential mating partner strains. Accordingly, higher secretors will be overall preferred by all strains of the opposite mating type. To detect pair-specific affinities, beyond such general effects in our analysis of mate choice (**Figure 3**), we compared the observed results to those expected under a null model. This null model considers the overall mating activity of each strain, which is influenced, among other factors, by its pheromone secretion level. Strains with high pheromone levels would thus be expected to mate across the board with multiple strains, but they might not raise above the null model expectation. Consequently, we were able to discern specific attraction and repulsion patterns between specific pairs of strains that go beyond potential general mating activity level differences.

Interestingly though, it is possible, that the coordination between pheromone levels and/or pheromone-to-receptor expression levels between partners may contribute to pair-specific mate choice. We found that genetically similar (though not identical) parents tend to attract each other (**Figure S8B**). An intriguing possibility is that genetic distance influences the coordination between pheromones or pheromone-to-receptor expression levels, thereby mediating mating affinity. Alternatively, genetic distance might impact other interactions crucial for successful mating, such as the interplay between agglutinins and sexual adhesins that facilitate parental fusion [58].

In this study, genetic distance was determined based on the overall number of SNPs that separate between any pair of strains, across the genome. However, a follow-up analysis could focus on genetic distance of specific mating-related genes. Such an examination may shed light on the mechanistic link between parental genetic distance and mate affinity.

Our findings demonstrate that yeast, particularly in glucose conditions, exhibits a capacity for adaptive mate affinity to enhance hybrids fitness (**Figure S8A**). This extends prior observations, including mate choice based on pheromone levels [47] or on cell size [24]. In our recent study, we established using the same system and strain collection, the heritability of fitness as a quantitative trait under the same growth conditions. We found that high fitness parents tend to yield higher fitness hybrids in glucose, but not in glycerol (Strauss *et al.* [51]). Here, we show that in glycerol, where offspring fitness is not directly correlated with parental fitness, there is no clear preference to mate with high-fitness partners (**Figure S10**). We also detect a correlation between heterozygosity level of the natural strain isolates and the inclination of parents derived from them toward outcrossing (**Figure S9**).

We note that our initial selection of a sub-set of ∼100 strains from the entire 1,011 strain collection, was designed to give a preference to strains with low heterozygosity levels [51]. As we now hypothesize that parental heterozygosity might dictate a higher level of outcrossing, our initial choice of strain may have resulted in an inclination to mate with genetically similar partners. This inclination thus reduces from the potential advantages associated with mating with high genetic distance partners in glycerol. The observed discrepancy suggests a nuanced conflict between the intrinsic inclination to outcross and the constraints imposed by the low heterozygosity of the original strains within our collection.

Comparing ’Forced’ mating and ’With-choice’ mating conditions (**Figure 2**), we observe better hybrids fitness in the ’With-choice’, condition especially in glycerol. Notably, the fitness assessment in the ’Forced’ and ’With-choice’ experiments is based on few generations of growth until reaching the stationary phase, while fitness assignments to variants are derived from extended competition experiments. Thus, pair-specific mating affinity in glycerol may optimize different fitness components compared to those measured in pooled competitions. This could explain the lack of observed enrichment in the over-represented mating group with high-fitness hybrids in glycerol.

In parallel to animals and plants, where mate selection has been observed [1]–[8], [11]–[18], our findings highlight the existence of pair-specific mating affinity in *S. cerevisiae*. This underscores the potential evolutionary significance of mate selection in sexual reproduction. A compelling question arises regarding the timing of the emergence of mate selection in evolution—whether it predates or arises concurrently with the ability for sexual mating. If a primordial mating system (such as yeast’s) lacks mate selection capabilities, the advantages of mating may be realized even without such specific mate affinity. Conversely, if mate affinity appeared simultaneously with sexual mating, it suggests that sexual reproduction can confer significantly greater benefits if accompanied by mate affinity and selection. Our results affirm that pair-specific mating affinity is integral to the sexual life of *S. cerevisiae*.

Two pivotal inquiries warrant further exploration: first, what cellular and molecular mechanisms underlie mate affinity in yeast; and second, why mate affinity in some conditions, e.g. glycerol here, does not optimize offspring fitness. Our observations prompt consideration on whether mate affinity in some conditions optimizes another aspect of the reproductive process. These questions open avenues for future investigations into the intricate interplay between mating preferences and evolutionary fitness in yeast.

## Materials and Methods

### Strains and media

Strains in this project were made based on the 1,011 natural isolates collection [40]. Chosen strains as well as their genetic engineering are presented at Strauss *et al* [51].

The following media types were used:

**SD Glu–** 6.7g/L yeast nitrogen base, 1.5g/L amino acid mix and 2% Glucose

**SD Gly–** 6.7g/L yeast nitrogen base, 1.5g/L amino acid mix and 2% Glycerol

**YPD -** 10g/L yeast extract, 20g/L peptone, 20g/L glucose

Antibiotic concentrations and initials:

Hygromycine B (Hyg) - 300mg/L,

Nourseothricin (NAT) – 100mg/L,

Kanamycin (G418) – 200mg/L

Zeocin (Zeo) – 200 mg/L

### Forced mating

In the “Forced” mating scheme we position in each well in a 96-well plate a pair of strains, one from the Mat**a** and one of Matα mating types. To perform pairwise mating, all verified haploid strains were taken out from the -80°C into YPD media with corresponding antibiotic (Hyg or NAT for Mat**a** or Matα respectively) in a 96-well plate using pinners. Strains were grown overnight (30°C, with shaking) and then diluted 1:1000 (SD-Glu) or 1:20 (SD-Gly) for another overnight incubation in 30°C to reach mid-log phase. OD was measured by plate reader (infinite 200, Tecan). Strains were diluted to reach equal OD. 40 96-well plates were filled with 100ul of either SD-Glu or SD-Gly with the addition of doxycycline (final concentration 10ug/ml). Then, 4ul (∼5E5 cells) from each Matα was inoculated into each well, using a multi-channel. Next, 4ul (∼5E5 cells) of each Mat**a** strain was inoculated into the same wells, resulting in ∼4000 pairwise mating reactions. Plates were incubated for 20 hours in 25°C. 10ul from each well was then collected into a reservoir to create the “Forced” mating pool.

### En masse mating

All verified haploid strains were taken out from the -80°C into YPD media with corresponding antibiotic (Hyg or NAT for Mat**a** or Matα respectively) in a 96-well plate using pinners. Strains were grown overnight (30°C, while shaking) and then diluted 1:1000 for another overnight incubation in 30°C. Strains were diluted 1:50 into either SD-Glu or SD-Gly and grown for another couple of hours in 30°C to reach mid-log phase. OD was measured by plate reader (infinite 200, Tecan). All Mat**a** strains and all Matα strains were mixed (separately for Mat**a** and Matα), based on the measured OD, such that they will have equal representation in the mix. Mixes were centrifuged and re-suspended in 0.2X volume to create a 5-fold increase in cell concentration (∼1E8 cells/ml). *En masse* mating was conducted by mixing 50ul (∼5E6 cells total) of each mating type mix in a 1ml medium (either SD-Glu or SD-Gly) with doxycycline (final concentration of 10ug/ml). Mating was executed for 20 hours in 25°C without shaking. One condition was done where strains were diluted 1:50 into SD-Glu and grown to reach mid-log, but then switched to SD-Gly after centrifugation, thus mating took place in SD-Gly. Three repetitions of *en masse* mating were done per media.

### Mating assay

Mating was performed as described in *en masse* mating section. Mating was done in three conditions; cells growth and mating on SD-Glu, cells growth and mating on SD-Gly or cells growth on SD-Glu followed by mating in SD-Gly (termed Glu switch to Gly experiment). Following mating, diploids were sorted based on their fluorescence signals as described below.

Sorting experiments were done in three consecutive days. Mat**a** and Matα mixes were kept in 4°C and used for mating in all three days. After 20 hours of mating, all mating reactions of the same media type were combined, and EDTA was added for a final concentration of 5mM.

Flow cytometry analysis and sorting were performed on a FACSAria Fusion instrument (BD Biosciences) equipped with a 405, 488, 561 and 640 nm lasers, using a 100 mm nozzle, controlled by BD FACS Diva software v8.0.1 (BD Biosciences), at The Weizmann Institute of Science Flow Cytometry Core Facility. Further analysis was performed using FlowJo software v10.2 (Tree Star). Cells were gated according to FSC and SSC to avoid debris and big aggregates. Another gate of high GFP and high mCherry was determined for sorting of diploid hybrids only.

Each day, the order of experiments to be sorted was changed to avoid bias. Cultures were kept in 4°C before and while sorting of other experiments took place.

Approximately 5E7 cells were sorted per experiment, resulting in ∼1E6 hybrids cells.

### Library preparation, sequencing and read analysis

For all experiments, library preparation was done using a similar protocol. Primer and DNA extraction methodology used in the different experiments vary, and are specified in Table 1.

**Table 1.**
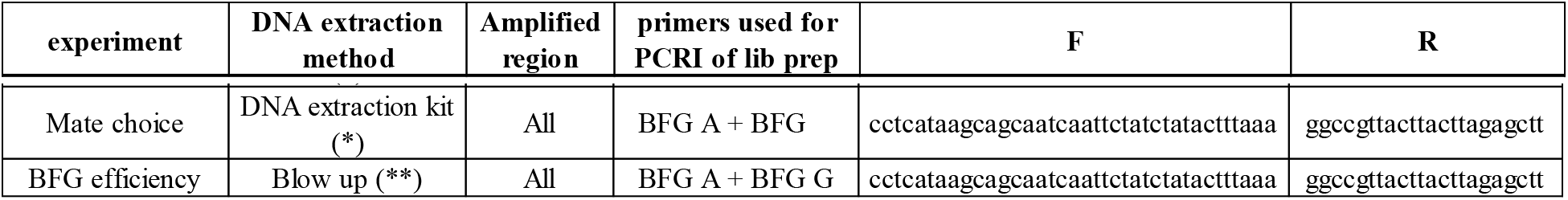
DNA extraction and library preparation protocols for each of the experiments. DNA extraction method and primers used in each of the described experiments. (*) MasterPure Yeast DNA Purification Kit by epicenter, (**) Lysis (Blow up) in 20mM NaOH and boiling for 15 minutes

Libraries for sequencing the barcode region were constructed by designing Plate-Row-Column PCR methodology; in which a first PCR is done using primers targeting the barcode region, plate barcode and tails that match Illumina adapters (F: ACGACGCTCTTCCGATCT**NNNNN***BFGprimer*, R: AGACGTGTGCTCTTCCGATCT**NNNNN***BFGprimer*

Capital letters corresponds to Illumina adaptors, N correspond to plate index and BFGprimer is the primer shown in Table 1(. First PCR was done in 25ul final volume with 2ul of template DNA (either genomic DNA or after cell lysis in 20mM NaOH and boiling for 15 minutes). PCR program: Tm of 60°C, elongation of 10 seconds, ∼20 cycles. Usually, 4 PCR reactions were done per experiments and they were pooled together after the first PCR (to avoid PCR biases). 2ul of the first PCR was used as a template for the second PCR.

A second PCR with the following primers

(F: AATGATACGGCGACCACCGAGATCTACACTCTTTCCCTACACGACGCTCTTCCGATCT, R:CAAGCAGAAGACGGCATACGAGAT***NNNNNNNN***GTGACTGGAGTTCAGACGTGTGCTCTTCCGATCT.

***N*** corresponds to Illumina index for library multiplexing) was carried out to attach the adapters for the Illumina run. PCR was done in 25ul volume. PCR program: Tm of 62°C, elongation of 10 seconds, ∼20 cycles. After second PCR libraries were cleaned using AMPure XP beads (Beckman Coulter, A63881) (∼1.2X ratio) to eliminate unspecific bands and primer dimers.

Amplicons were sequenced using paired-end methodology, on the NovaSeq platform (Illumina). All PCR reactions were done using KAPA HiFi HotStart ReadyMix (Roch, KK2601 ).

After initial de-multiplexing by the Illumina platform, libraries were further separated based on the plate index using cutadapt [59]. All reads were further processed by cutadapt to leave only the barcode region. For alignment, a synthetic genome from all strains’ barcodes was made using bowtie2 (bowtie2-build command). Alignment was performed using bowtie2 as well. Read counts per variant were determined by in-house script.

### BFG efficiency

Chosen strains (**Table S1**) were used for measurement and calculation of BFG efficiency. Strains were taken out from -80°C and grown at 30°C overnight in YPD while shaking. Cells were diluted 1:1000 and grown at 30°C overnight in SD-Glu while shaking. OD was measured using plate reader (infinite 200, Tecan), and all strains were diluted to equal OD values. Matα strains (∼1E7 cells) were inoculated into the corresponding row of 96-well plates while that Mat**a** strains (∼1E7 cells) were added to the columns of each plate and allowed to mate for 20 hours at 25°C without shaking in SD-Glu supplemented with 10ug/ml Doxycycline. After 20 hours mating, cells were diluted 1:120 into fresh SD-Glu supplemented with 10ug/ml Doxycycline, Hyg and NAT, and continued growing for one day. Cells were then diluted 1:120 into SD-Glu + Hyg + NAT (without Doxycycline) and grown for one day. After 5 days of daily dilutions 1:120 (∼30 generations), plates were FACS analyzed to determine the hybrid ratio in each culture. Hybrids were scored using GFP/mCherry gating. In most wells, hybrids were found to reach a proportion of over 90% of the culture. One row and one column had no hybrids, indicating a problem with that strain (data not shown). At this point, DNA was extracted by boiling the cells in 20mM NaOH for 15 minutes and library were constructed with the most upstream and downstream primers (**Figure S1B** primers A + G, and **Table1** in Materials & Methods). BFG efficiency was calculated by dividing the number of reads with fused barcodes with number of reads with original barcodes for each pair.

### Pairwise mating for determining mating efficiency

Relevant strains (see **Table S2**) were taken from -80°C and were grown for 48 hours at 30°C in YPD while shaking. Cells were diluted 1:1000 into SD-Glu or SD-Gly and incubated overnight in 30°C while shaking. Cells were diluted 1:50 and allowed growing for a couple of hours to reach mid-log. OD was measured using a plate reader (infinity 200, Tecan). Two strains (one Mat**a** and one Matα strain) were inoculated into the same well such that their cell number is equal (∼1E7 cells). Mating was executed for 20 hours in 25°C without shaking. All mating reactions were done in 3 repetitions.

Mating efficiency for each pair of strains was determined using FACS (Attune) using 96-well plate module. Cultures were diluted 1:200 before FACS into SD with 5mM EDTA. Each culture was divided into 3 populations based on GFP/mCherry ratio; high GFP and low mCherry were considered Mat**a** cells, high mCherry and low GFP were the Matα cells, and high mCherry and GFP are the hybrids. Mating efficiency (ME) was calculated based on the following formula: *ME* = #*hybrids*/*min*(*Mat**a***, *Mat⍺*. Mating efficiency was calculated as the number of hybrids divided by the minimum of Mat**a** and Matα number of cells.

## Acknowledgements

We thank the Pilpel lab for extensive feedback and discussions. We thank Naama Barkai for useful comments. We thank Frederick Roth lab and Dayag Sheykhkarimli from Toronto university for sharing the BFG system with us, and for advices in the cloning process.

All cloning, but adding the BFG into the construct, was done by Yoav Peled and the Structural Proteomics unit in Weizmann Institute of Science. FACS sorting was done with Tomer Meir Salame in the FACS unit at the Weizmann Institute of Science. Sequencing was performed at the Crown Institute of Genomics, G-INCPM at Weizmann Institute of Science.

We thank the Israel Science Foundation, the Yeda-Sela Foundation at the Weizmann Institute, and the Minerva Foundation for grant support. YP is the Ben-May Professorial Chair and a Kimmel Investigator.

