## Supplementary note for "Adaptive Mating Among Natural Strains of *S. cerevisiae*"

### Supplementary note - Considering additional factors that may affect mating affinity determination

### BFG calculations

##### BFG efficiency vary between strains, but can be predicted per pair by the parents’ average BFG efficiency

In order to detect each hybrid, yeast were transformed with a sophisticated system enabling the fusion of barcodes initially found on different chromosomes (in this case, the two parental chromosomes). Thus, the barcode fusion (BFG) efficiency is an essential part of detecting pairs as deduced from hybrid offspring’s frequency. To calculate BFG efficiency, 16 Mat**a** strains and *9* Matα strains were chosen **(Table S1**) and pairwise mating between all of them (1*44* pairs) was done. After pairwise mating, hybrids were selected by double antibiotics and sequenced to find BFG efficiency (see Materials & Methods, BFG efficiency part for more details). BFG efficiency values varied across different pairs of strains (**Figure S4A**). In addition, the average BFG efficiency of strains (calculated over all partners with whom it was crossed) also varies (**Figure S4B**).

These results indicate that the ability to induce the recombinase and or complete the fusion event differ between different genetic backgrounds. Since BFG is instrumental for the ability to identify the identity of hybrids we wanted to verify whether this efficiency can be estimated for all possible pairs. For that, an expected BFG efficiency was calculated per pair by multiplying the average efficiency of each partner. The correlation between expected efficiency and calculated efficiency is very high, and significant (**Figure S4C**), although not entirely on the x=y line**.** This result suggests that BFG efficiency is not a unique property of pair of strains, but rather unique for each haploid strain, and that the hybrid BFG efficiency is solely a simple multiplication between the parents. Given the strong correlation, we concluded that BFG efficiency can be predicted for all possible pairs with high accuracy. To validate that the BFG efficiency does not bias the results of mating affinity, we plotted pairs BFG efficiency in either the over-represented, as expected or absent mate affinity groups. By and large, no significant differences are found (**Figure S6**).

#### Pairwise mating efficiency calculations

##### Mating efficiency vary between strains, but can be predicted per pair by the parents’ average mating efficiency

16 Mat**a** strains and 15 Matα strains were used to test pairwise Mating Efficiency (ME) (**Table S2**). After pairwise mating between all strains, wells were analyzed by FACS, and ratio of offspring were measured based on fluorescence (Mat**a** is labeled with GFP, Matα is labeled with RFP, thus offspring are double positive). We then used this measurement to calculate the ME between each two strains. Since no choice is given in this experiment, this value represents the maximal/actual mating ability between the two parents in question.

ME was defined as $ME = double positive cells/(double positive cells+ min(Mat\mathbf{a}, Mat\alpha)$).

**Figure S5A** shows that ME varies between pairs. **Figure S5B and S5C** shows that measured ME is correlated to the calculated ME of a pair of strains i.e to the multiplication of the average ME of Mat**a** and Matα parents (parental mean ME was calculated by averaging all wells that a specific strain).

The correlation is high in glucose, but low in glycerol. For further validation that the ME doesn’t bias the results of mating affinity, we have plotted the average ME of offspring from either the over-represented, as expected or absent mate affinity groups. No significance different was seen in any of the two carbon sources (**Figure S7**).

#### Fitness variance as a potential bias for mating affinity

A possible concern is that the differences between hybrids frequency observed following mating (**Figure 3B**) may reflect not only mating affinity but also variations in fitness between strains during the 20 hours of the mating reaction. These fitness differences could result from either the uneven growth of parents, leading to the over-representation of specific parents, or variations in the immediate growth of offspring post-mating.

While we cannot completely rule out residual effects of fitness differences, we contend that this factor alone is unlikely to fully explain the mate choice results presented in **Figure 3B**. It is crucial to emphasize that the *en masse* mating reactions were conducted in a manner designed to minimize yeast growth, specifically aiming to eliminate the influence of fitness differences. First, a high number of parental strains (with carefully accounted even representation of each parent) were added to each reaction (~10E7 cells per ml), ensuring that even under optimal growth conditions, cells could undergo only 3-4 generations before reaching the stationary phase. Second, mating took place under conditions far from optimal for yeast growth, including relatively low temperature and the absence of shaking.

Note that the observed difference in hybrids frequency after mating between the "as-expected" and "over-represented" groups exceeded three orders of magnitude (**Figure 3B**). Even under the assumption of no growth of some hybrids (or parents), the maximum frequency difference achievable with 4 generations due to fitness differences alone is approximately 16-fold. Thus, the observed hybrids frequency cannot be attributed to fitness differences in either the parents or offspring, indicating a genuine mating affinity.
