## Supplementary figures for "Adaptive Mating Among Natural Strains of *S. cerevisiae*"

### Slide 1
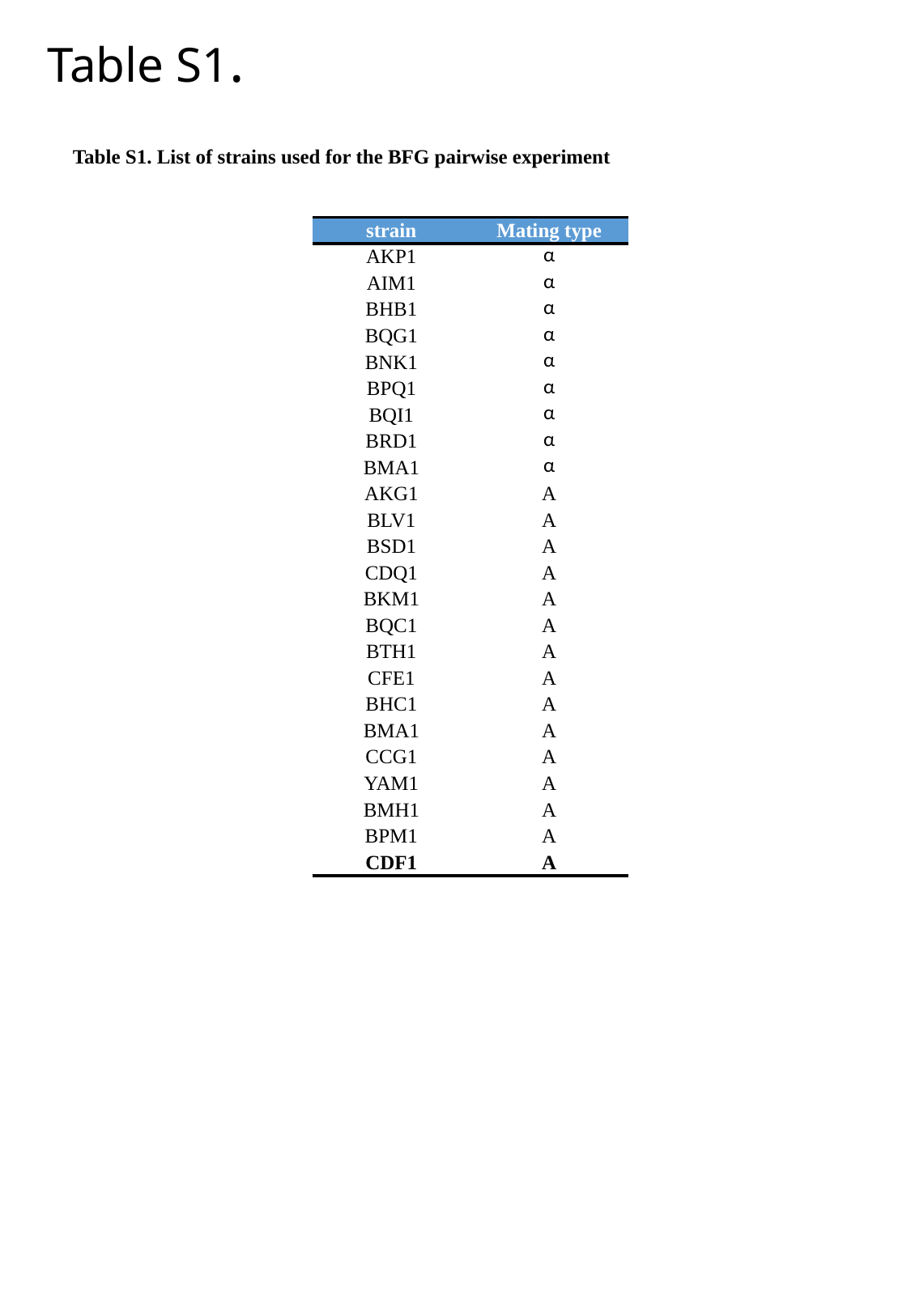

Table S1.
Table S1. List of strains used for the BFG pairwise experiment
| strain | Mating type |
| --- | --- |
| AKP1 | α |
| AIM1 | α |
| BHB1 | α |
| BQG1 | α |
| BNK1 | α |
| BPQ1 | α |
| BQI1 | α |
| BRD1 | α |
| BMA1 | α |
| AKG1 | A |
| BLV1 | A |
| BSD1 | A |
| CDQ1 | A |
| BKM1 | A |
| BQC1 | A |
| BTH1 | A |
| CFE1 | A |
| BHC1 | A |
| BMA1 | A |
| CCG1 | A |
| YAM1 | A |
| BMH1 | A |
| BPM1 | A |
| CDF1 | A |

### Slide 2
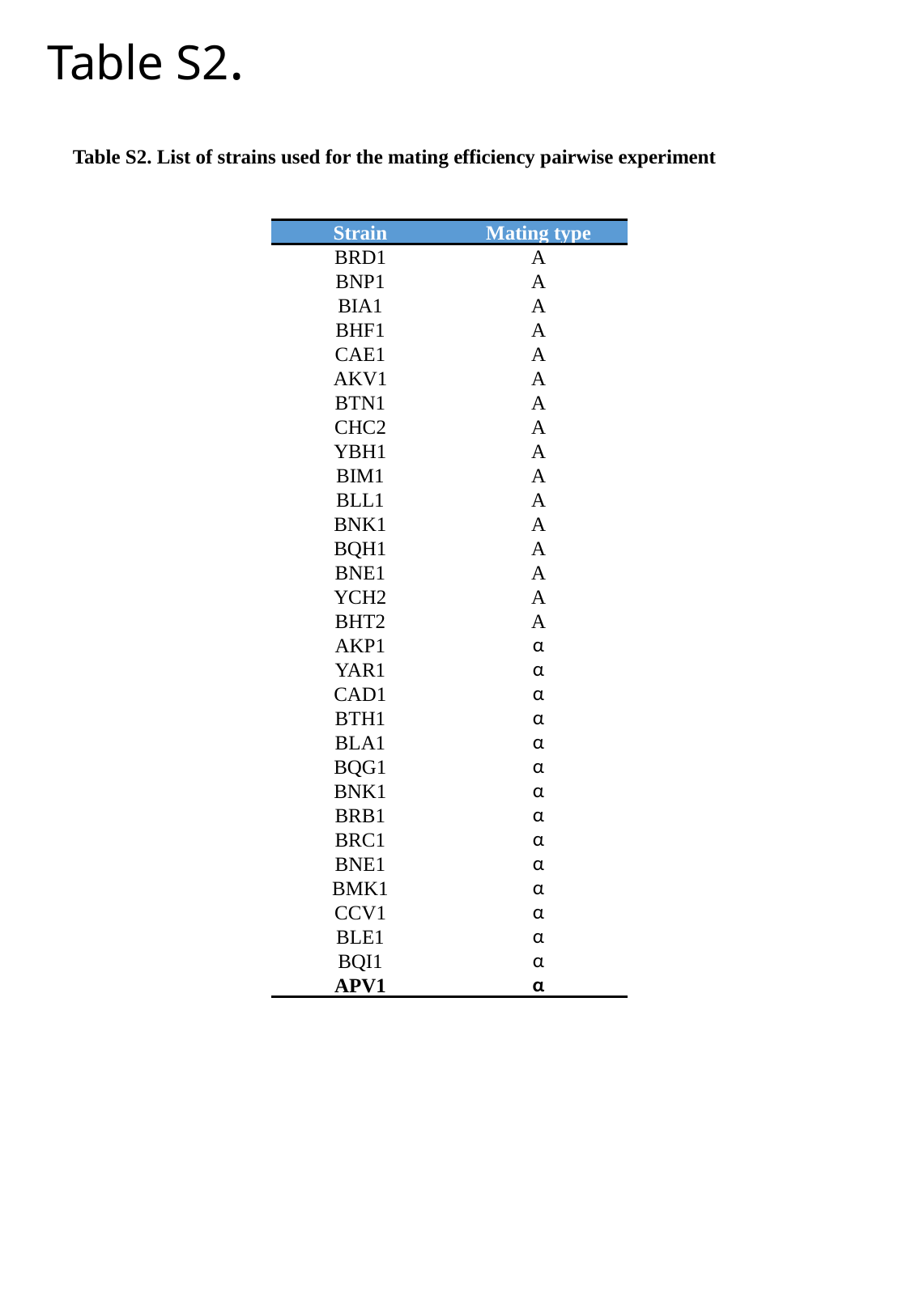

Table S2.
Table S2. List of strains used for the mating efficiency pairwise experiment
| Strain | Mating type |
| --- | --- |
| BRD1 | A |
| BNP1 | A |
| BIA1 | A |
| BHF1 | A |
| CAE1 | A |
| AKV1 | A |
| BTN1 | A |
| CHC2 | A |
| YBH1 | A |
| BIM1 | A |
| BLL1 | A |
| BNK1 | A |
| BQH1 | A |
| BNE1 | A |
| YCH2 | A |
| BHT2 | A |
| AKP1 | α |
| YAR1 | α |
| CAD1 | α |
| BTH1 | α |
| BLA1 | α |
| BQG1 | α |
| BNK1 | α |
| BRB1 | α |
| BRC1 | α |
| BNE1 | α |
| BMK1 | α |
| CCV1 | α |
| BLE1 | α |
| BQI1 | α |
| APV1 | α |

### Slide 3
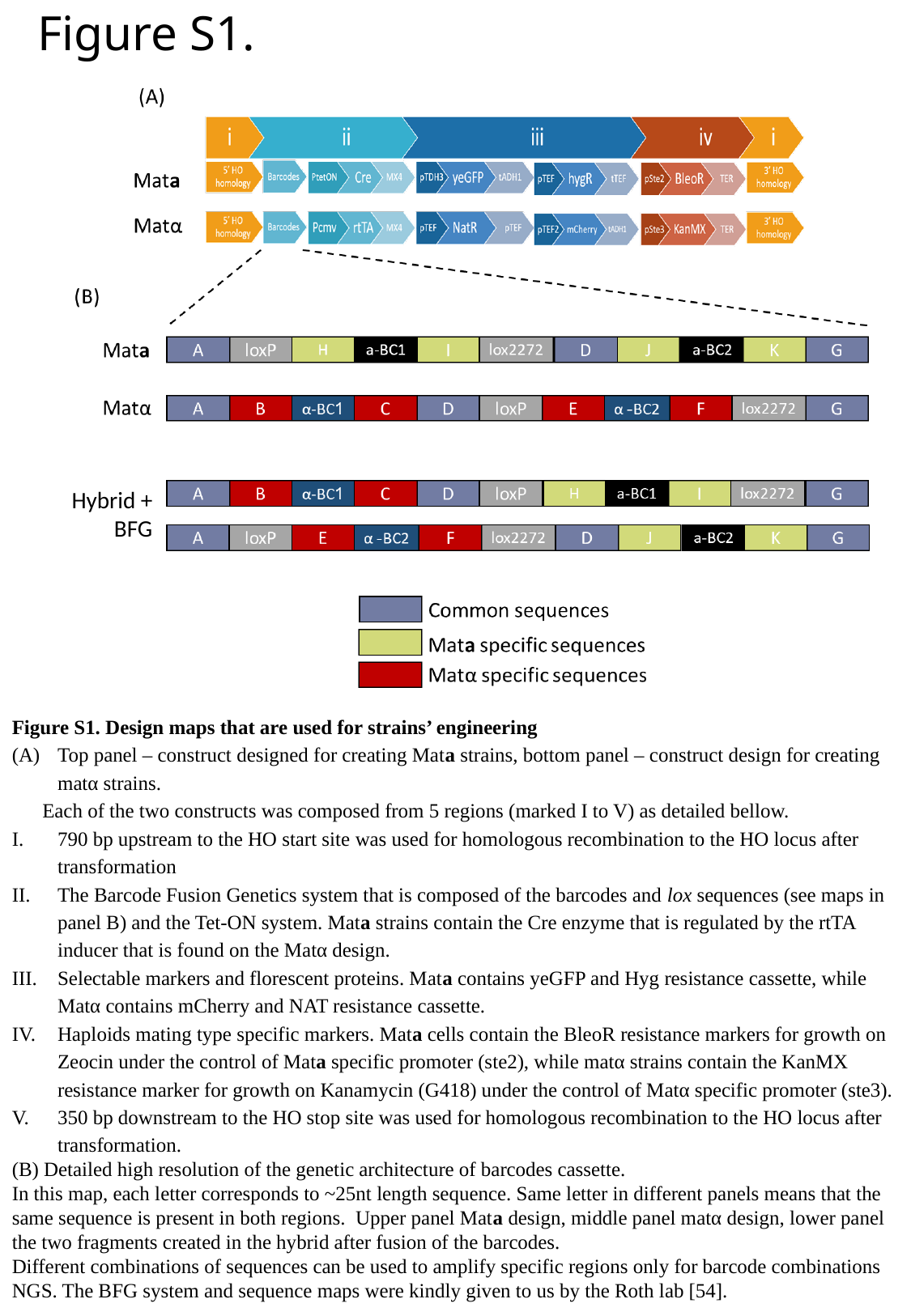

#### Figure S1.
Hybrid + BFG
Figure S1. Design maps that are used for strains’ engineering
Top panel – construct designed for creating Mata strains, bottom panel – construct design for creating matα strains.
Each of the two constructs was composed from 5 regions (marked I to V) as detailed bellow.
790 bp upstream to the HO start site was used for homologous recombination to the HO locus after transformation
The Barcode Fusion Genetics system that is composed of the barcodes and lox sequences (see maps in panel B) and the Tet-ON system. Mata strains contain the Cre enzyme that is regulated by the rtTA inducer that is found on the Matα design.
Selectable markers and florescent proteins. Mata contains yeGFP and Hyg resistance cassette, while Matα contains mCherry and NAT resistance cassette.
Haploids mating type specific markers. Mata cells contain the BleoR resistance markers for growth on Zeocin under the control of Mata specific promoter (ste2), while matα strains contain the KanMX resistance marker for growth on Kanamycin (G418) under the control of Matα specific promoter (ste3).
350 bp downstream to the HO stop site was used for homologous recombination to the HO locus after transformation.
(B) Detailed high resolution of the genetic architecture of barcodes cassette.
In this map, each letter corresponds to ~25nt length sequence. Same letter in different panels means that the same sequence is present in both regions. Upper panel Mata design, middle panel matα design, lower panel the two fragments created in the hybrid after fusion of the barcodes.
Different combinations of sequences can be used to amplify specific regions only for barcode combinations NGS. The BFG system and sequence maps were kindly given to us by the Roth lab [54].

### Slide 4
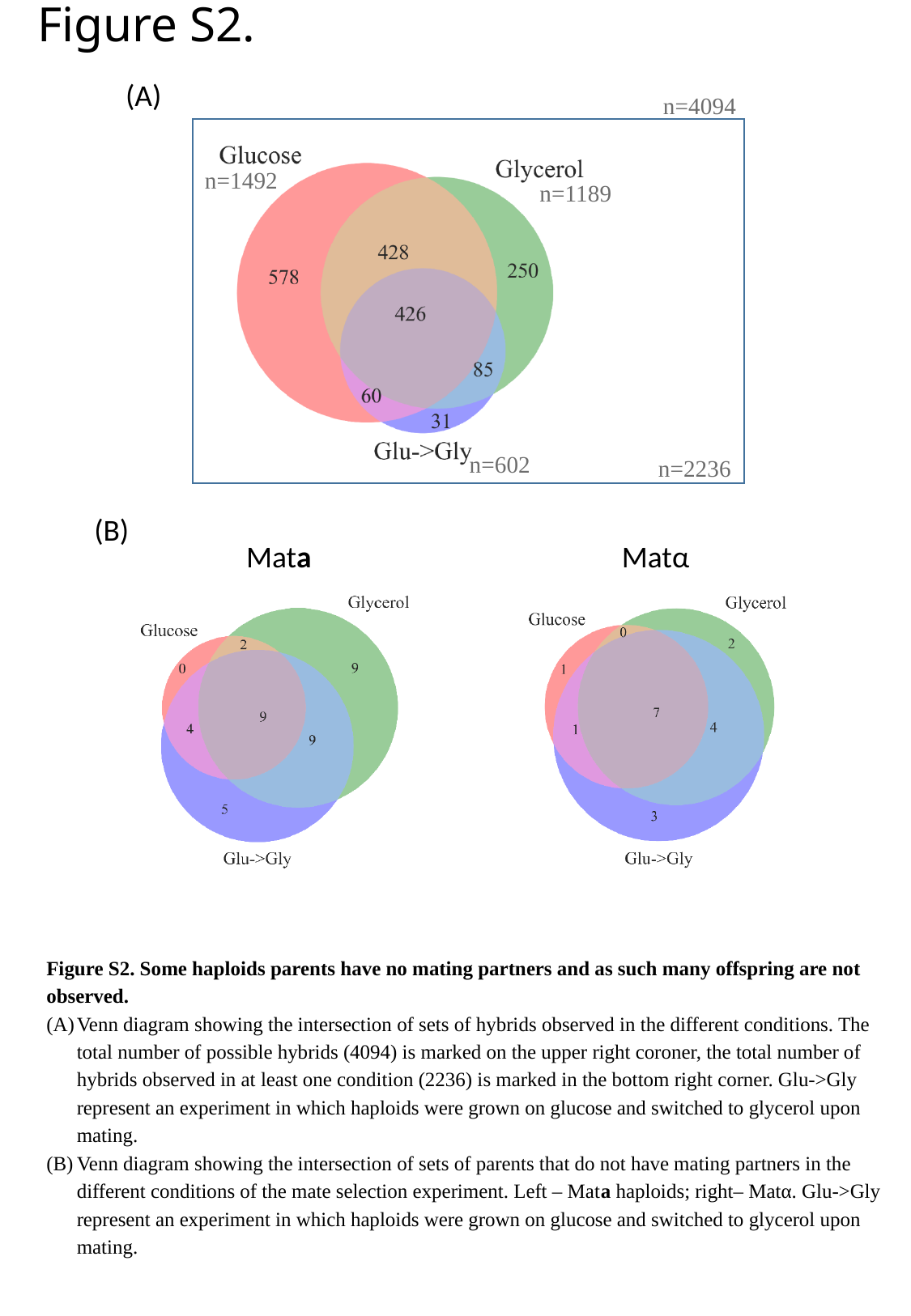

Figure S2.
(A)
n=4094
n=1492
n=1189
n=602
n=2236
(B)
Mata
Matα
Figure S2. Some haploids parents have no mating partners and as such many offspring are not observed.
Venn diagram showing the intersection of sets of hybrids observed in the different conditions. The total number of possible hybrids (4094) is marked on the upper right coroner, the total number of hybrids observed in at least one condition (2236) is marked in the bottom right corner. Glu->Gly represent an experiment in which haploids were grown on glucose and switched to glycerol upon mating.
Venn diagram showing the intersection of sets of parents that do not have mating partners in the different conditions of the mate selection experiment. Left – Mata haploids; right– Matα. Glu->Gly represent an experiment in which haploids were grown on glucose and switched to glycerol upon mating.

### Slide 5
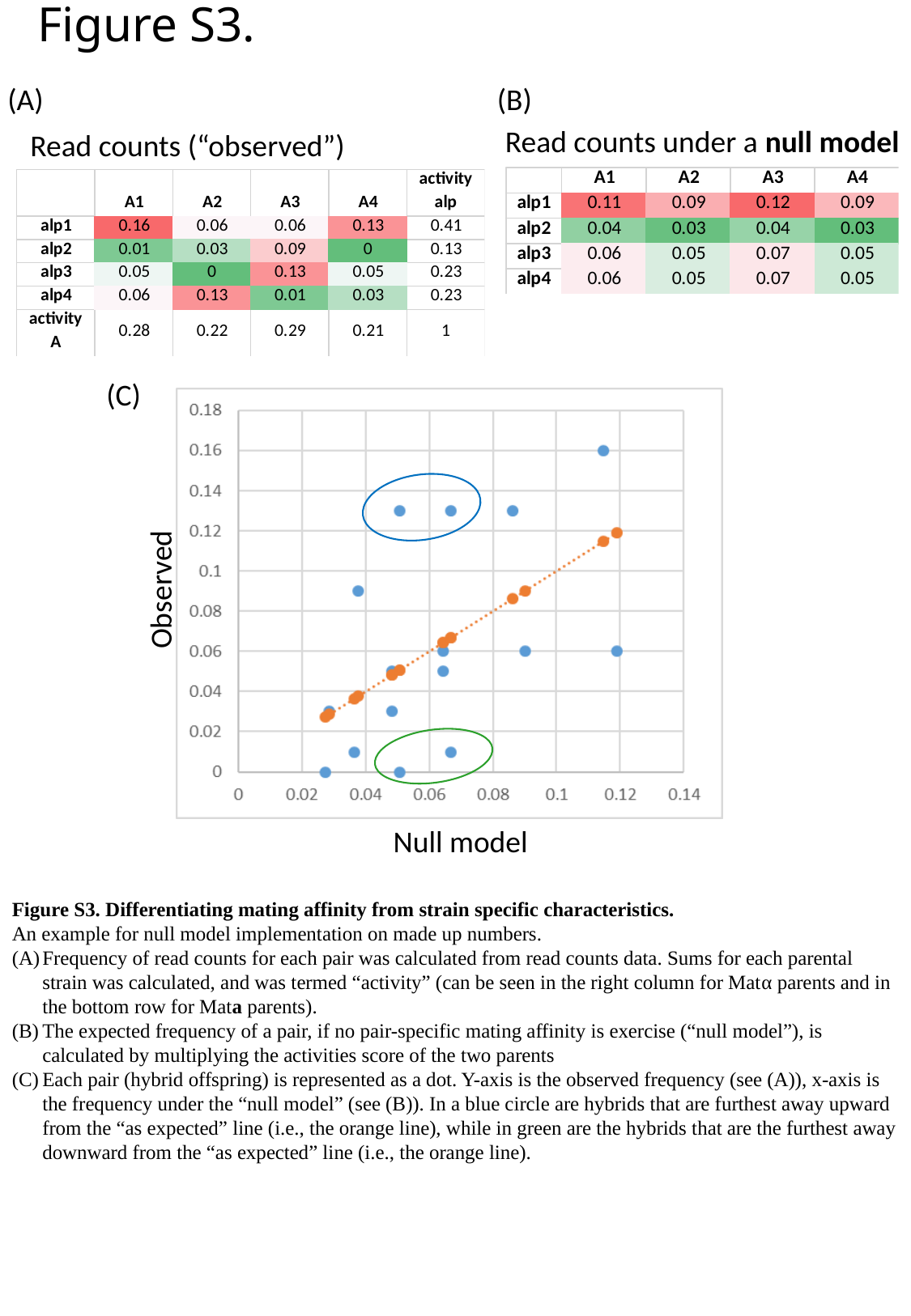

Figure S3.
(A)
(B)
Read counts under a null model
Read counts (“observed”)
(C)
Observed
Null model
Figure S3. Differentiating mating affinity from strain specific characteristics.
An example for null model implementation on made up numbers.
Frequency of read counts for each pair was calculated from read counts data. Sums for each parental strain was calculated, and was termed “activity” (can be seen in the right column for Matα parents and in the bottom row for Mata parents).
The expected frequency of a pair, if no pair-specific mating affinity is exercise (“null model”), is calculated by multiplying the activities score of the two parents
Each pair (hybrid offspring) is represented as a dot. Y-axis is the observed frequency (see (A)), x-axis is the frequency under the “null model” (see (B)). In a blue circle are hybrids that are furthest away upward from the “as expected” line (i.e., the orange line), while in green are the hybrids that are the furthest away downward from the “as expected” line (i.e., the orange line).

### Slide 6
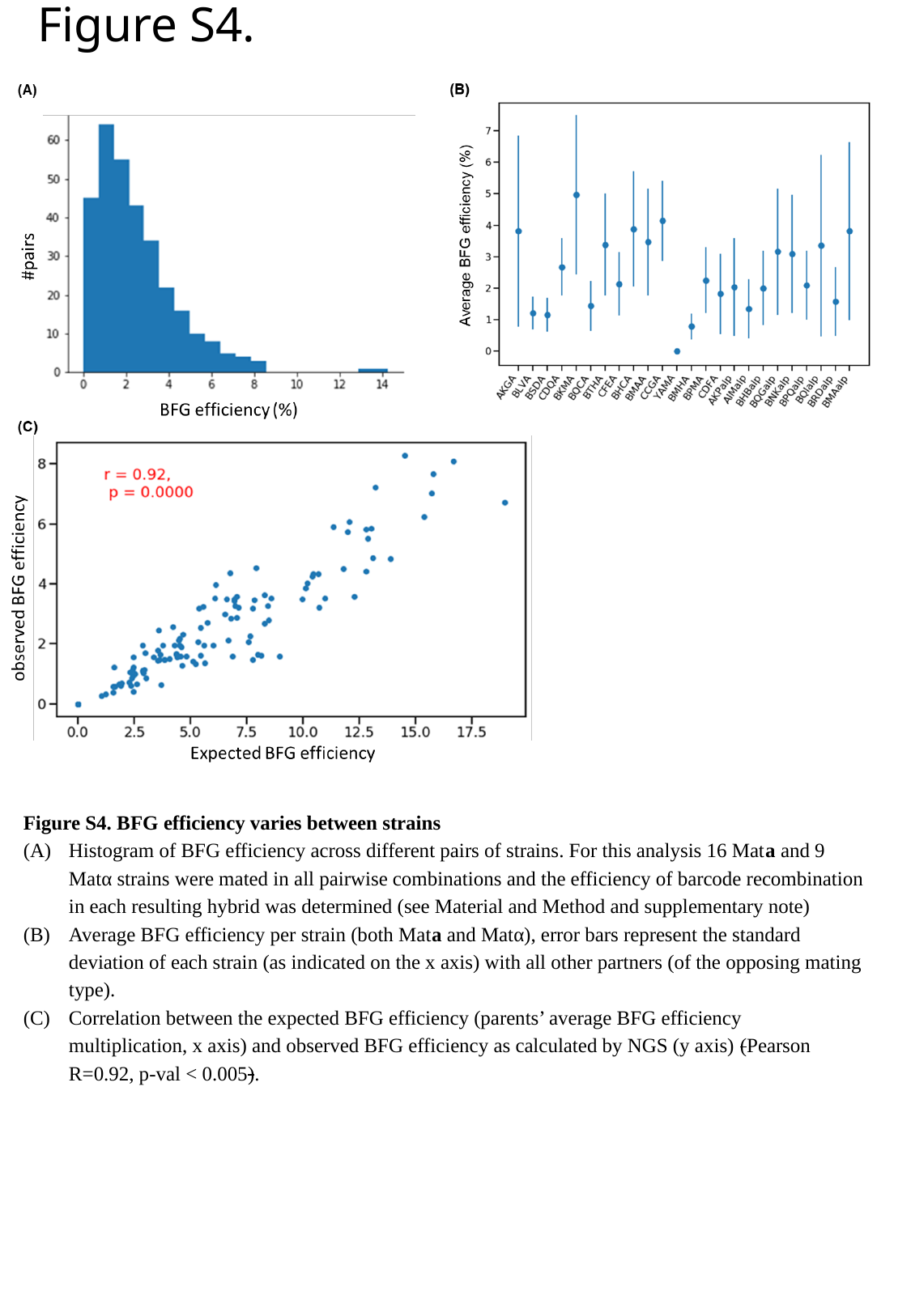

Figure S4.
Figure S4. BFG efficiency varies between strains
Histogram of BFG efficiency across different pairs of strains. For this analysis 16 Mata and 9 Matα strains were mated in all pairwise combinations and the efficiency of barcode recombination in each resulting hybrid was determined (see Material and Method and supplementary note)
Average BFG efficiency per strain (both Mata and Matα), error bars represent the standard deviation of each strain (as indicated on the x axis) with all other partners (of the opposing mating type).
Correlation between the expected BFG efficiency (parents’ average BFG efficiency multiplication, x axis) and observed BFG efficiency as calculated by NGS (y axis) (Pearson R=0.92, p-val < 0.005).

### Slide 7
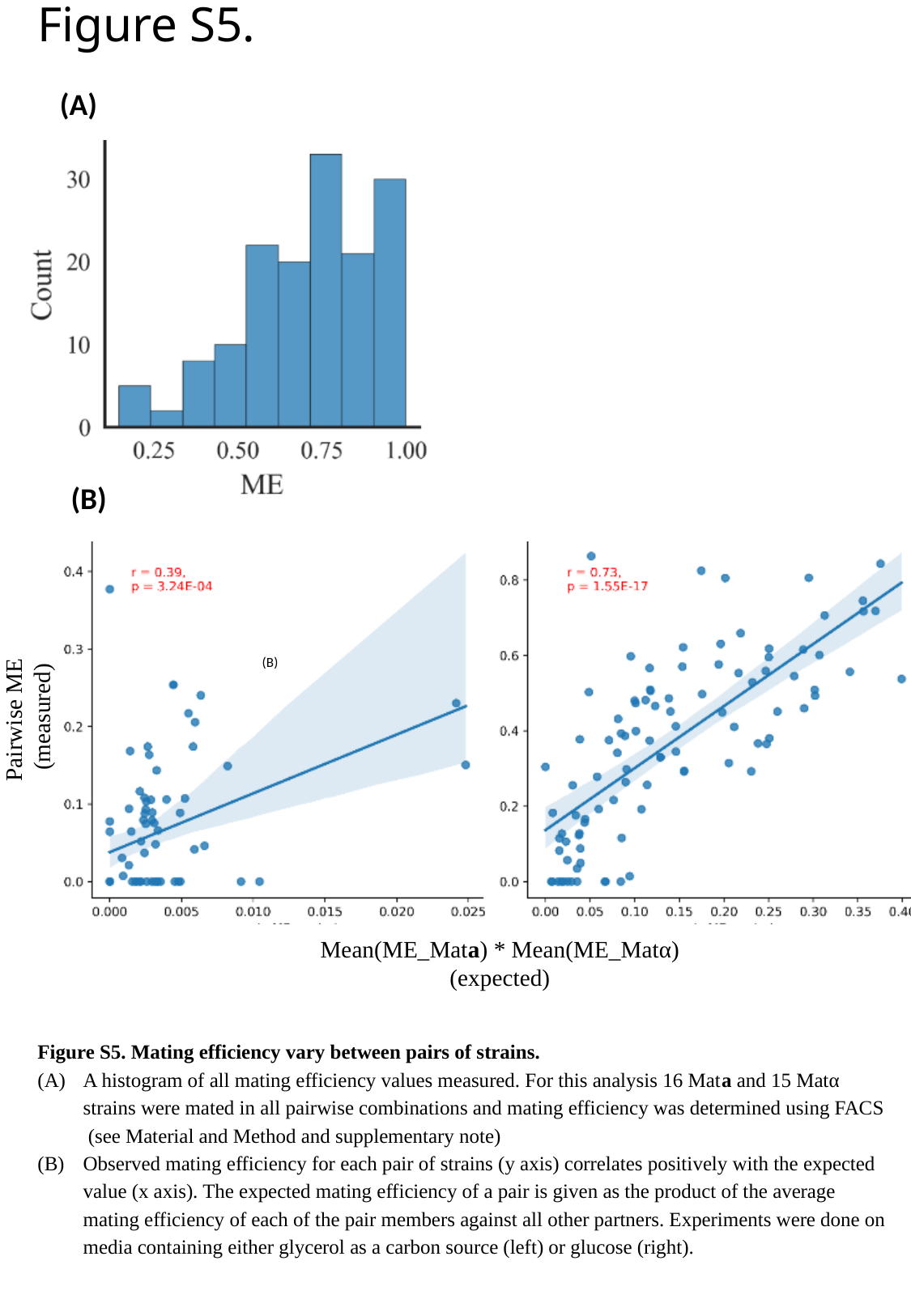

Figure S5.
(A)
(B)
(B)
(C)
Pairwise ME
 (measured)
Mean(ME_Mata) * Mean(ME_Matα)
(expected)
Figure S5. Mating efficiency vary between pairs of strains.
A histogram of all mating efficiency values measured. For this analysis 16 Mata and 15 Matα strains were mated in all pairwise combinations and mating efficiency was determined using FACS (see Material and Method and supplementary note)
Observed mating efficiency for each pair of strains (y axis) correlates positively with the expected value (x axis). The expected mating efficiency of a pair is given as the product of the average mating efficiency of each of the pair members against all other partners. Experiments were done on media containing either glycerol as a carbon source (left) or glucose (right).

### Slide 8
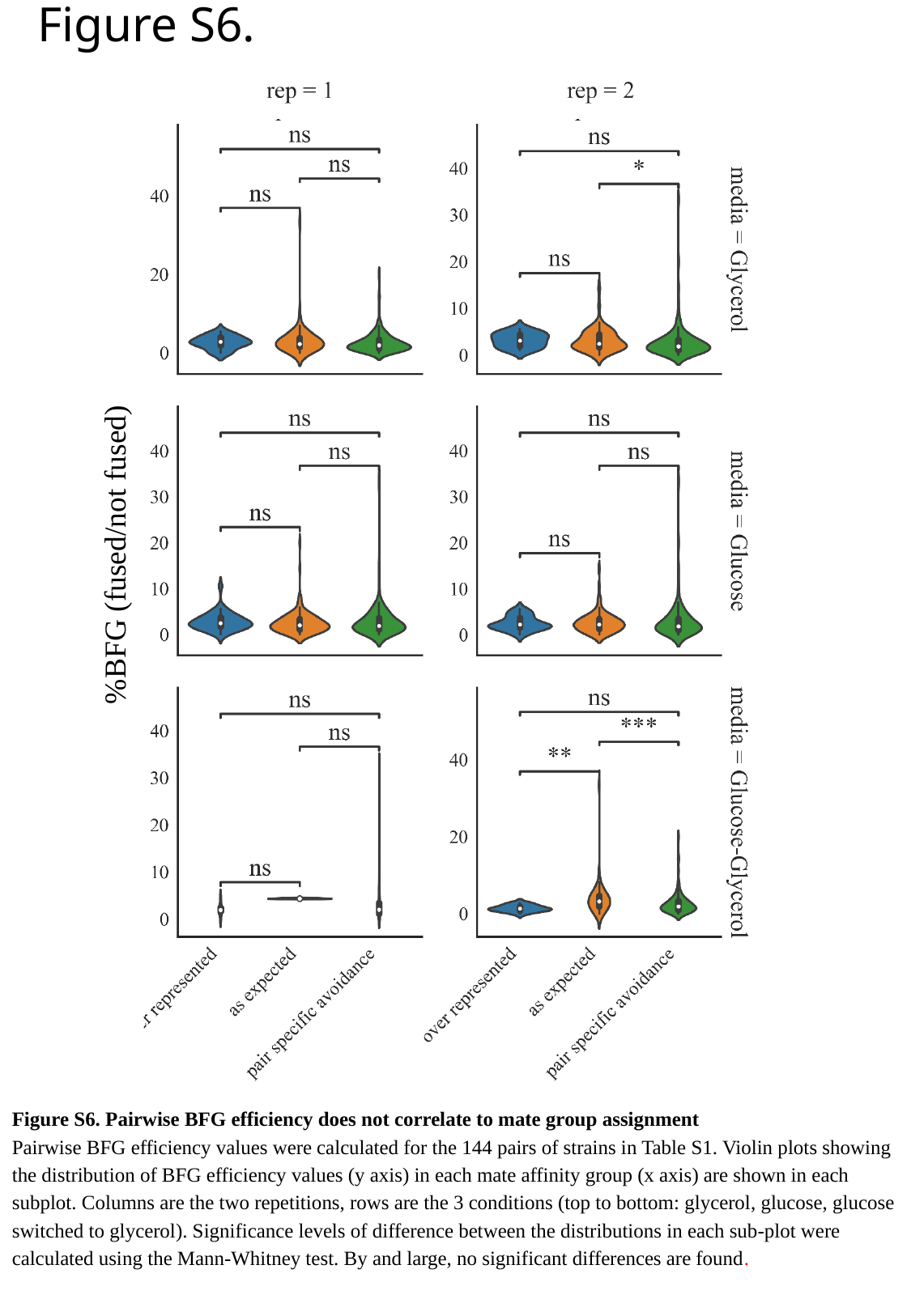

Figure S6.
%BFG (fused/not fused)
Figure S6. Pairwise BFG efficiency does not correlate to mate group assignment
Pairwise BFG efficiency values were calculated for the 144 pairs of strains in Table S1. Violin plots showing the distribution of BFG efficiency values (y axis) in each mate affinity group (x axis) are shown in each subplot. Columns are the two repetitions, rows are the 3 conditions (top to bottom: glycerol, glucose, glucose switched to glycerol). Significance levels of difference between the distributions in each sub-plot were calculated using the Mann-Whitney test. By and large, no significant differences are found.

### Slide 9
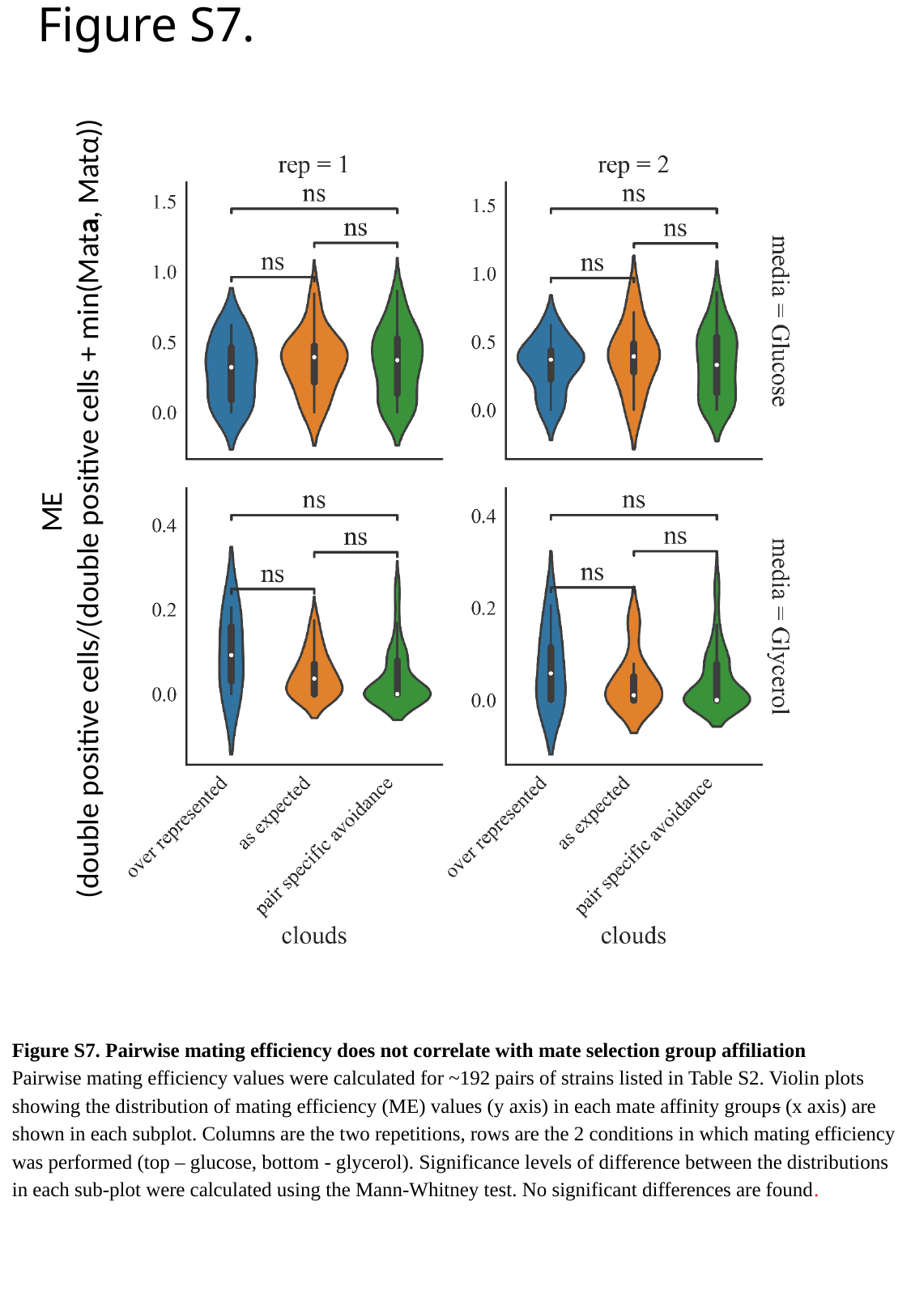

Figure S7.
ME
 (double positive cells/(double positive cells + min(Mata, Matα))
Figure S7. Pairwise mating efficiency does not correlate with mate selection group affiliation
Pairwise mating efficiency values were calculated for ~192 pairs of strains listed in Table S2. Violin plots showing the distribution of mating efficiency (ME) values (y axis) in each mate affinity groups (x axis) are shown in each subplot. Columns are the two repetitions, rows are the 2 conditions in which mating efficiency was performed (top – glucose, bottom - glycerol). Significance levels of difference between the distributions in each sub-plot were calculated using the Mann-Whitney test. No significant differences are found.

### Slide 10
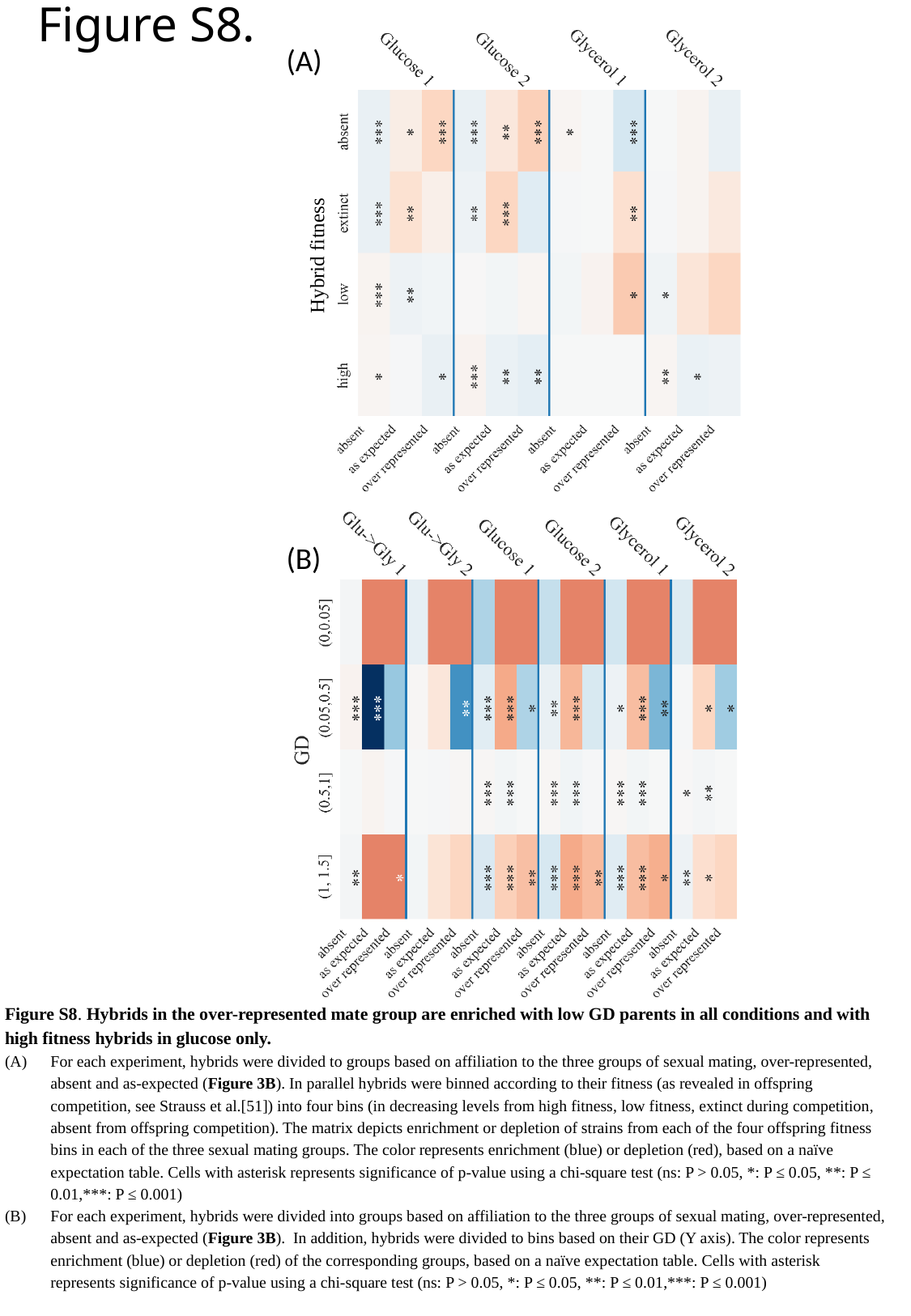

Figure S8.
(A)
Hybrid fitness
(B)
Figure S8. Hybrids in the over-represented mate group are enriched with low GD parents in all conditions and with high fitness hybrids in glucose only.
For each experiment, hybrids were divided to groups based on affiliation to the three groups of sexual mating, over-represented, absent and as-expected (Figure 3B). In parallel hybrids were binned according to their fitness (as revealed in offspring competition, see Strauss et al.[51]) into four bins (in decreasing levels from high fitness, low fitness, extinct during competition, absent from offspring competition). The matrix depicts enrichment or depletion of strains from each of the four offspring fitness bins in each of the three sexual mating groups. The color represents enrichment (blue) or depletion (red), based on a naïve expectation table. Cells with asterisk represents significance of p-value using a chi-square test (ns: P > 0.05, *: P ≤ 0.05, **: P ≤ 0.01,***: P ≤ 0.001)
For each experiment, hybrids were divided into groups based on affiliation to the three groups of sexual mating, over-represented, absent and as-expected (Figure 3B). In addition, hybrids were divided to bins based on their GD (Y axis). The color represents enrichment (blue) or depletion (red) of the corresponding groups, based on a naïve expectation table. Cells with asterisk represents significance of p-value using a chi-square test (ns: P > 0.05, *: P ≤ 0.05, **: P ≤ 0.01,***: P ≤ 0.001)

### Slide 11
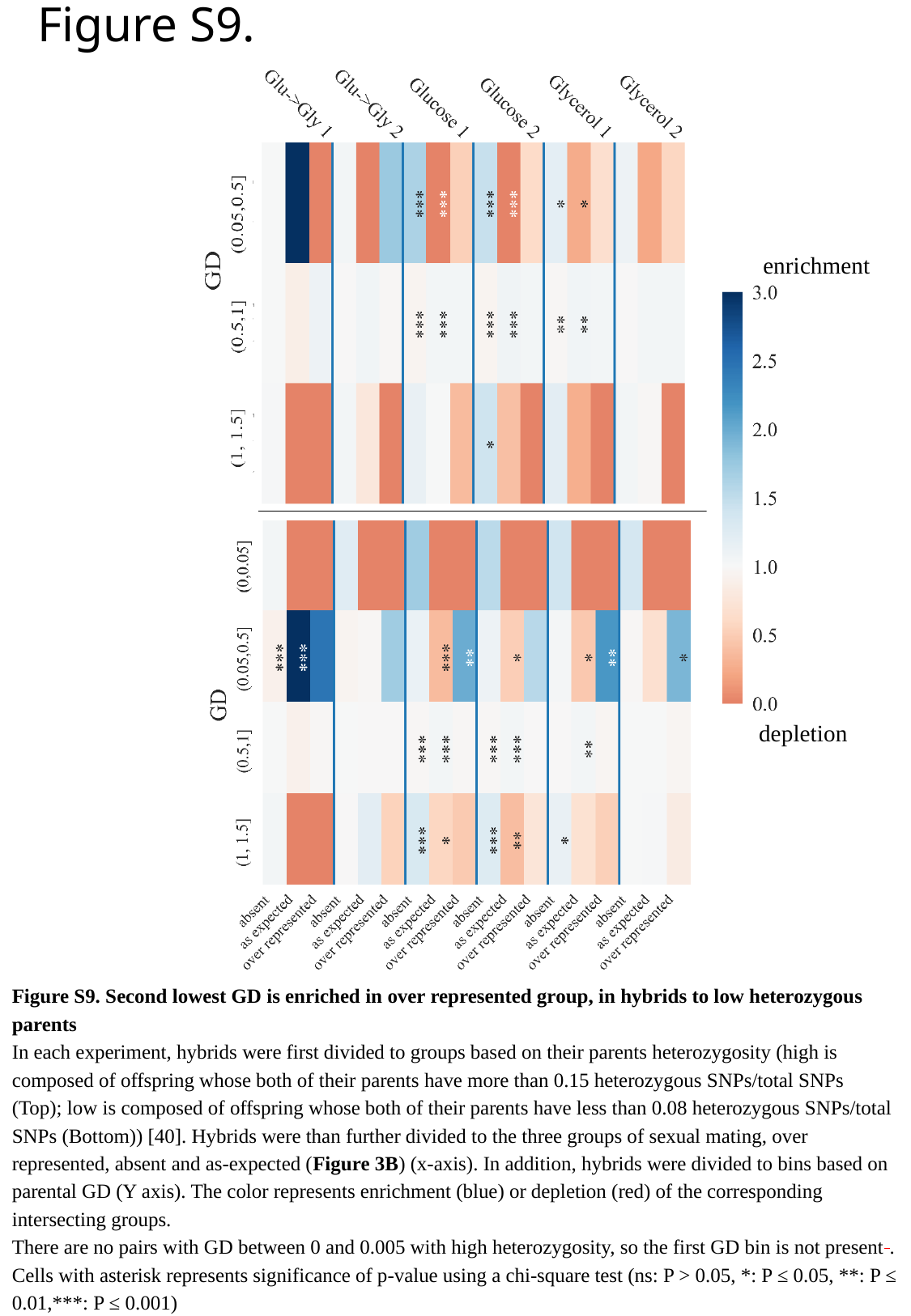

Figure S9.
enrichment
depletion
Figure S9. Second lowest GD is enriched in over represented group, in hybrids to low heterozygous parents
In each experiment, hybrids were first divided to groups based on their parents heterozygosity (high is composed of offspring whose both of their parents have more than 0.15 heterozygous SNPs/total SNPs (Top); low is composed of offspring whose both of their parents have less than 0.08 heterozygous SNPs/total SNPs (Bottom)) [40]. Hybrids were than further divided to the three groups of sexual mating, over represented, absent and as-expected (Figure 3B) (x-axis). In addition, hybrids were divided to bins based on parental GD (Y axis). The color represents enrichment (blue) or depletion (red) of the corresponding intersecting groups.
There are no pairs with GD between 0 and 0.005 with high heterozygosity, so the first GD bin is not present .
Cells with asterisk represents significance of p-value using a chi-square test (ns: P > 0.05, *: P ≤ 0.05, **: P ≤ 0.01,***: P ≤ 0.001)

### Slide 12
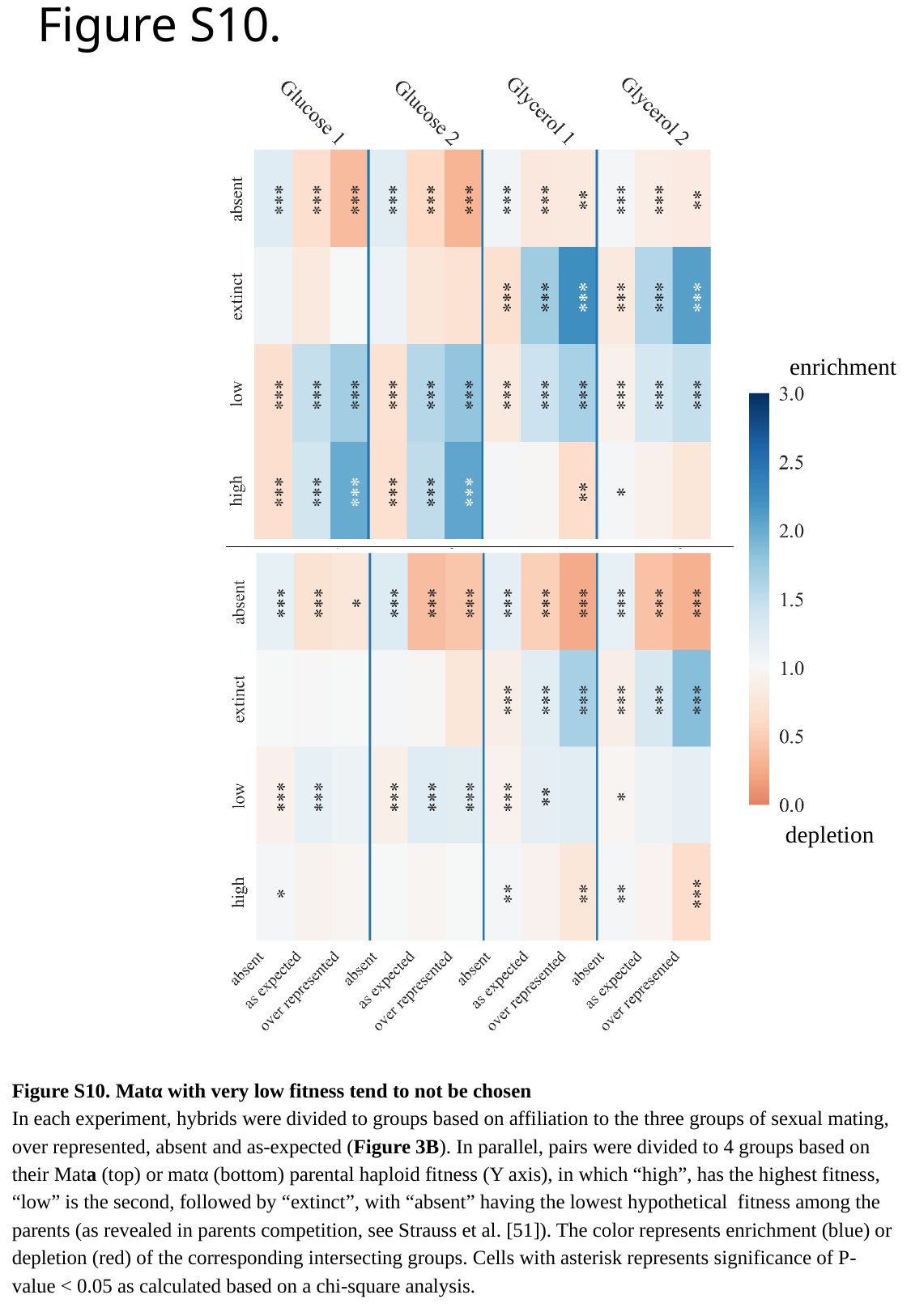

Figure S10.
enrichment
depletion
Figure S10. Matα with very low fitness tend to not be chosen
In each experiment, hybrids were divided to groups based on affiliation to the three groups of sexual mating, over represented, absent and as-expected (Figure 3B). In parallel, pairs were divided to 4 groups based on their Mata (top) or matα (bottom) parental haploid fitness (Y axis), in which “high”, has the highest fitness, “low” is the second, followed by “extinct”, with “absent” having the lowest hypothetical fitness among the parents (as revealed in parents competition, see Strauss et al. [51]). The color represents enrichment (blue) or depletion (red) of the corresponding intersecting groups. Cells with asterisk represents significance of P-value < 0.05 as calculated based on a chi-square analysis.
